# Divergent roles of SOX2 in human and mouse germ cell specification related to X-linked gene dosage effects

**DOI:** 10.1101/2024.06.25.599839

**Authors:** Wenteng He, Qing Luo, Jian Zhao, Mengting Wang, Luohua Feng, Allan Zhao, Ahmed Reda, Eva Lindgren, Jan-Bernd Strukenborg, Jiayu Chen, Qiaolin Deng

**Author notes:** These authors contributed equally (Wenteng He, Qing Luo).

## Abstract

Human primordial germ cell-like cells (hPGCLCs) can be generated from pluripotent stem cells (PSCs) but the differentiation efficiency of female hPSCs is often lower than that of male hPSCs. Moreover, Klinefelter Syndrome (KS), a condition characterized by an extra X-chromosome in males, often presents the failure of germline specification and infertility. In this study, we investigate how X-linked gene dosage affects hPGCLCs specification potential in both healthy and diseased conditions. We reveal that the X-chromosome plays a multifaceted role in modulating hPGCLCs induction. The inhibitory effects on TGF-beta/Activin A and BMP pathways by escape genes IGSF1 and CHRDL1, respectively, are demonstrated by the increased yield of hPGCLCs with knockdown experiments. Importantly, our results identified the intriguing role of SOX2 that is upregulated by the escape gene *USP9X* in hPGCLCs specification, highlighting a species-specific difference from the mouse model. The elevated *USP9X*-*SOX2* regulatory axis profoundly influences cellular metabolism, mitochondrial morphology, and progenitor competence, thereby affecting hPGCLCs induction. Furthermore, the inability to downregulate SOX2 and upregulate SOX17 in response to BMP signaling impedes downstream gene activation due to motif binding competition. These findings shed novel insights into the hPGC specification by elucidating the differential roles of SOX2 versus SOX17 between mice and humans, influenced by X-linked gene dosage effects. Additionally, our results offer potential applications for improving the induction and survival efficiency of hPGCLCs from hPSCs, facilitating disease modeling and mechanistic studies.

**Highlights:**

- Downregulation of three X-linked genes, i.e. IGSF1, CHRDL1 and USP9X, enhanced the differentiation efficiency of hPGCLCs
- SOX2 as a downstream of human-specific escape gene USP9X plays a multifacet role against hPGCLCs specification
- Failure to timely downregulate SOX2 and upregulate SOX17 interferes downstream gene activation likely due to motif binding competition

## Introduction

Reconstitution of functional gametes *in vitro* from human pluripotent stem cells (hPSCs) stands for one of the most challenging goals in the stem cell research field because of the intricated genetic and epigenetic reprogramming during the germline specification.^1^ An increasing understanding of the molecular events during germ cell development has provided the blueprint to realize this goal. Concurrently, human induced pluripotent stem cells (hiPSCs) derived from various diseases have proven to be valuable tools for elucidating gene functions and exploring epigenetic regulations associated with cell-type differentiation.^2^ Therefore, we are now equipped to more comprehensively understand the mechanisms underlying infertility.

In mammals, females undergo X-chromosome inactivation (XCI) during early development to balance the dosage of X-linked gene expression between males and females.^3^ While the majority of genes from the inactivated X-chromosome are simultaneously silenced, certain genes evade XCI and are expressed on both X-chromosomes, known as escapees.^4^ In humans, 15∼%25 of X-linked genes are known escapees, in which some genes consistently evade inactivation in all tissues (i.e., constitutive escapees), while others are context-specific (i.e., facultative escapees).^5, 6^ In comparison, only 3–7% of X-linked genes escape XCI in mice.^7^ Several studies, including ours, have shown that differentiation kinetics differ between male and female PSCs, as the X-linked gene dosage affects the transcriptome, epigenome and pluripotency exit of these cells.^8–11^ Moreover, increasing evidence suggests that XCI can be eroded over time in culture due to unfavorable conditions, such as high lithium chloride in mTeSR1 medium,^12^ leading to increased expression of X-linked genes.^13^ Proteome analysis also revealed that XCI erosion has a broader downstream effect than what was previously thought,^14^ suggesting that X-linked gene dosages have cascaded effects through impacting autosomal genes. Furthermore, a large-scale analysis of female hPSCs lines found that eroded XCI is associated with poor differentiation,^15–18^ as cells also tend to maintain their previous XCI status. Notably, the differentiation efficiency of female hPSCs towards human primordial germ cell-like cells (hPGCLCs) is often lower than that of male hPSCs.^19, 20^ Also, X-linked gene dosage has significant implications for the pathology of Klinefelter Syndrome (KS), a condition characterized by the presence of at least one extra X chromosome in males (i.e., 47, XXY), which results in male infertility, autoimmune disease and metabolic disorders etc.^21^ These phenotypic manifestations of KS patients are associated with the extent of XCI.^17^ However, the underlying mechanisms by which X-linked gene dosage affects hPGCLC specification are poorly understood in both healthy and diseased conditions.

Here, we systematically investigated male and female hPSCs as well as two KS patients (47, XXY) derived iPSCs, i.e., KS1 and KS2, together with the control male iPSCs. Our results showed that increased X-linked gene dosage greatly impaired the efficiency of hPGCLCs specification. We discovered that knockdown of XCI escapees *CHRDL1* and *IGSF1*, shown to regulate activin A^22^ and BMP^22^ signaling, respectively, could greatly enhance hPGCLCs differentiation efficiency in KS1 but to a much lesser extent in KS2 or female hPSCs with erosion of XCI (i.e., HS980). In contrast, decreased expression of constitutive escapee *UPS9X* was found to promote hPGCLCs specification in both KS1 and KS2, as well as HS980. Moreover, we found that *SOX2*, as the downstream target of *UPS9X,* played an important role against hPGCLCs specification. Elevated *SOX2* expression led to compromised oxidative phosphorylation and increased mitochondrial clustering in KS and HS980. Additionally, it skewed fate transition towards ectoderm-like rather than mesoderm-like state, which is crucial for hPGCLCs induction. Furthermore, sustained expression of *SOX2* during early PGCLC induction caused extensive cell death upon activation of *SOX17* by BMP signaling as *SOX2* and *SOX17* exhibit mutual exclusivity in germ cell specification.

Our results suggested that SOX2 acts as a roadblock in human PGC induction mainly by interfering with cellular competence, mitochondrial function, and downstream gene regulation network. Since SOX2 is regulated by human specific XCI escapee *USP9X*, balancing X-linked gene dosage balance is more critical in human germline differentiation. These findings shed novel insights into the longstanding question of why SOX2 function is not conserved in mouse and human PGC development. Moreover, our findings could be leveraged to improve the differentiation efficiency and survival of hPGCLCs from iPSCs with suboptimal XCI states.

## Results

### Differentiation of hPGCLCs is influenced by the X-linked gene dosage

Earlier studies have highlighted that male hPSC lines exhibit a relatively higher efficiency in hPGCLC differentiation than their female counterparts.^19, 20^ Given that XCI often occurs in female hPSCs cultured in the primed pluripotent condition, we hypothesized that X-linked gene dosage imbalance due to escapees might underlie this sex-specific difference. Notably, KS patients suffer from azoospermia, and this severity is often associated with the XCI extent on the additional X chromosome,^17^ which further suggests that the X-linked gene dosage could affect the specification of human germ cells, although the underlying mechanism remains elusive. To comprehend this, we studied two KS patient-derived iPSC lines (KS1 and KS2) in addition to a male iPSC (XY) line that we previously characterized.^23^ Both KS1 and KS2 lines maintain proper pluripotency and can normally form all three germ layers.^23^ However, KS2 exhibits higher X-linked gene expression due to the erosion of XCI, allowing us to further explore the effects of X-linked gene dosage on the germline specification. For comparison, we also included two female lines H9 and H980, with HS980 showing the erosion of XCI.^23^

Differentiation of these hPSCs into hPGCLCs was conducted using a well-established two-step protocol via the incipient mesoderm-like cells (iMeLCs) followed by hPGCLCs induction for four days (D4) (Figure 1A).^24^ At the hPSCs stage, all lines displayed similar characteristic colony-like morphology with well-defined boundaries (Figure 1B). Interestingly, at the iMeLCs stage, XY iPSC and H9 lines showed typical flat epithelial features with clear cell-to-cell boundaries. In contrast, KS1, KS2 and HS980 lines showed various degrees of colony-like morphology, indicating a compromised differentiation (Figure 1B). Furthermore, at the D4 following hPGCLC induction, both KS lines exhibited significant cell death within the spheroids compared to the XY line, with KS2 showing particularly poor spheroid formation and lower cellular viability (Figures 1B and S1A-S1C). Similarly, for the female lines, HS980 displayed worse cell viability compared to H9 (Figures 1B and S1B-S1C). We further assessed hPGCLCs induction efficiency in the D4 hPGCLC spheroids by measuring double positivity of surface markers EpCAM and INTEGRIN-α6 (ITGA6). Consistent with spheroid morphology, the two KS lines and HS980 yielded below 1% hPGCLCs compared to approximately 24% and 12% hPGCLCs in XY and H9, respectively (Figures 1C and 1D). Consequently, XY and H9 obtained high hPGCLC efficiencies (Figure S1D).

**Figure 1.**
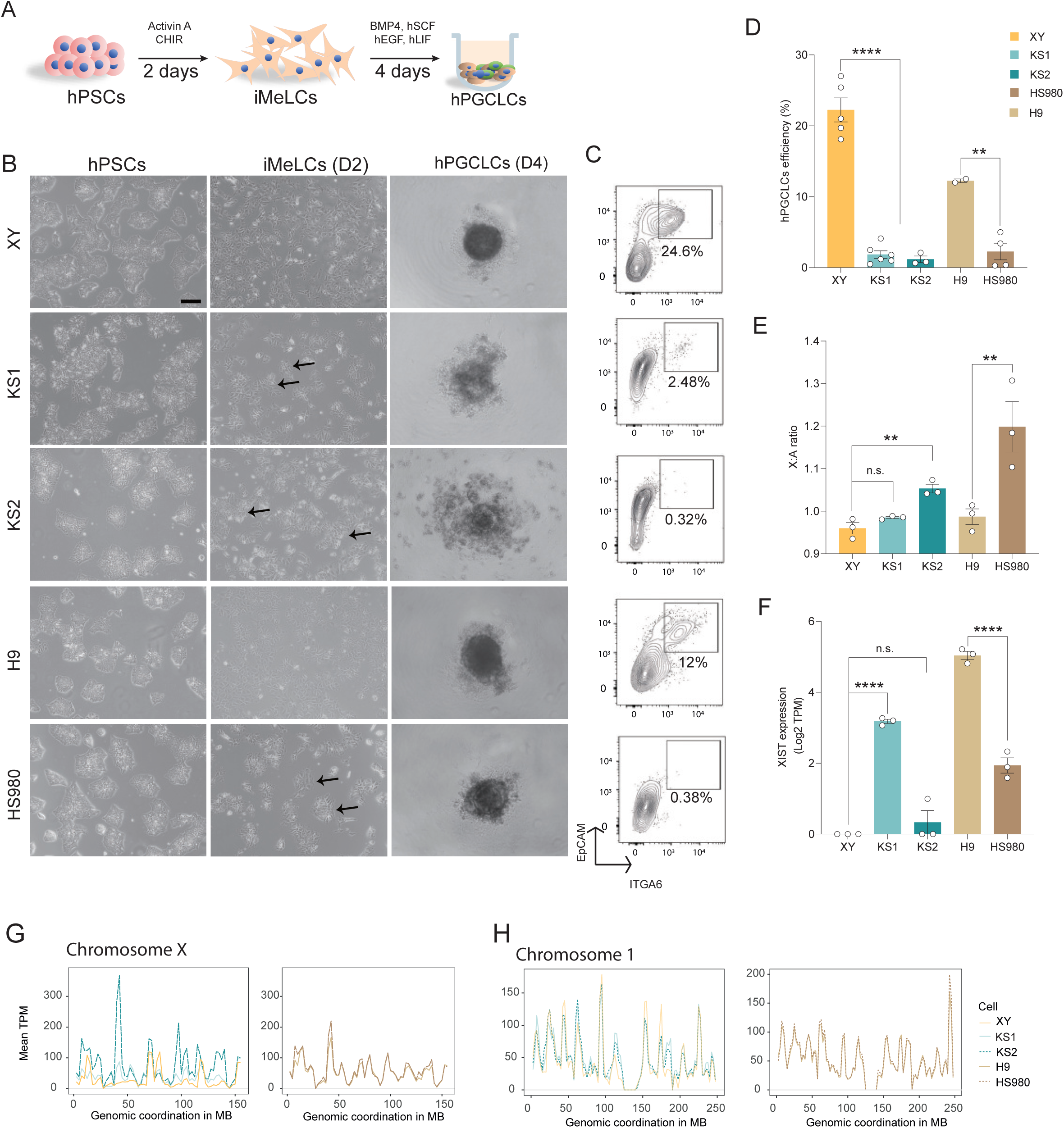
Differentiation of hPGCLCs is affected by the X-linked gene dosage. (A) Schematic illustration of the hPGCLC differentiation workflow. (B) Representative morphology of cultured PSC, iMeLC and D4 hPGCLC spheroid. Scale bar, 200 μm. Black arrow identifies colony-like iMeLCs. (C) D4 hPGCLC ratio identified by FACS with double markers of ITGA6 and EpCAM. (D) Statistics of the D4 hPGCLC efficiency. (E) X:A ratios of KS1, KS2, XY, H9 and HS980. (F) *XIST* expression of the corresponding cell lines in (E). (G) Gene expression levels shown on genomic coordination of the X chromosome of KS1, KS2, XY, H9 and HS980. (H) Gene expression levels shown on genomic coordination of Chromosome 1 of KS1, KS2, XY, H9 and HS980. Data in (D), (E), and (F) are shown as mean ± SEM. n.s., non-significant; \*\**p* < 0.01; \*\*\*\**p* < 0.0001 by one-way ANOVA comparison.

To investigate whether the reduced efficiencies were associated with X-linked gene dosage, we first analyzed the X chromosome to autosome (X:A) ratio across the five lines. We performed bulk RNA-seq for our three lines, XY, KS1 and KS2, and utilized published RNA-seq data for H9 and HS980. As expected, KS2 and HS980 exhibited significantly higher X:A ratios (Figure 1E). Moreover, KS2 showed minimal *XIST* and HS980 also showed low *XIST* expression (Figure 1F). These findings confirmed the erosion of XCI in KS2 and HS980, as previously reported. Through mapping the gene expressions onto chromosome coordinates, we found that KS2 had the highest expression output along the entire X-chromosome, followed by KS1 and XY in line with *XIST* expression (Figure 1G). The differences were less pronounced between H980 with H9, except for certain loci, aligning with *XIST* expression in HS980 (Figure 1G). Notably, the constitutive escape gene *USP9X* is in a region with peaked expression (Figure 1G). Reassuringly, we did not observe large chromosome-wide differences among all lines for individual autosomes (Figures 1H and S2). These results further supported the role of X-linked gene dosage in germ cell specification.

### Downregulation of escape genes increases the efficiency of hPGCLC specification

To further understand how X-linked genes affect two stages towards germ cell induction, we firstly compared the global transcriptomic profiles of our two KS lines and the control XY line with those of the published female line UCLA1 and male line UCLA2 at the PSC and iMeLC stages. UCLA1 and UCLA2 are two well-characterized hESCs capable of hPGCLCs specification. Notably, UCLA1, as the female line, exhibited low efficiency in hPGCLCs specification.^20^ For convenience, we designated them as UCLA-F (female) and UCLA-M (male) in our study. Principal component analysis (PCA) of the transcriptomic data revealed that the two KS cell lines were closely aligned with the XY control at both the iPSC and iMeLC stages (Figure 2A). Similarly, both UCLA lines were also clustered accordingly (Figure 2A). Notably, UCLA-F and UCLA-M showed the greatest differences at the iMeLC stage, likely correlating with the low yield of UCLA-F (Figure 2A). These findings suggested that aneuploidy in KS lines did not alter their PSC and iMeLC identities, corroborating our previous report that KS lines had a normal capacity for three germ layer specifications.^23^

**Figure 2.**
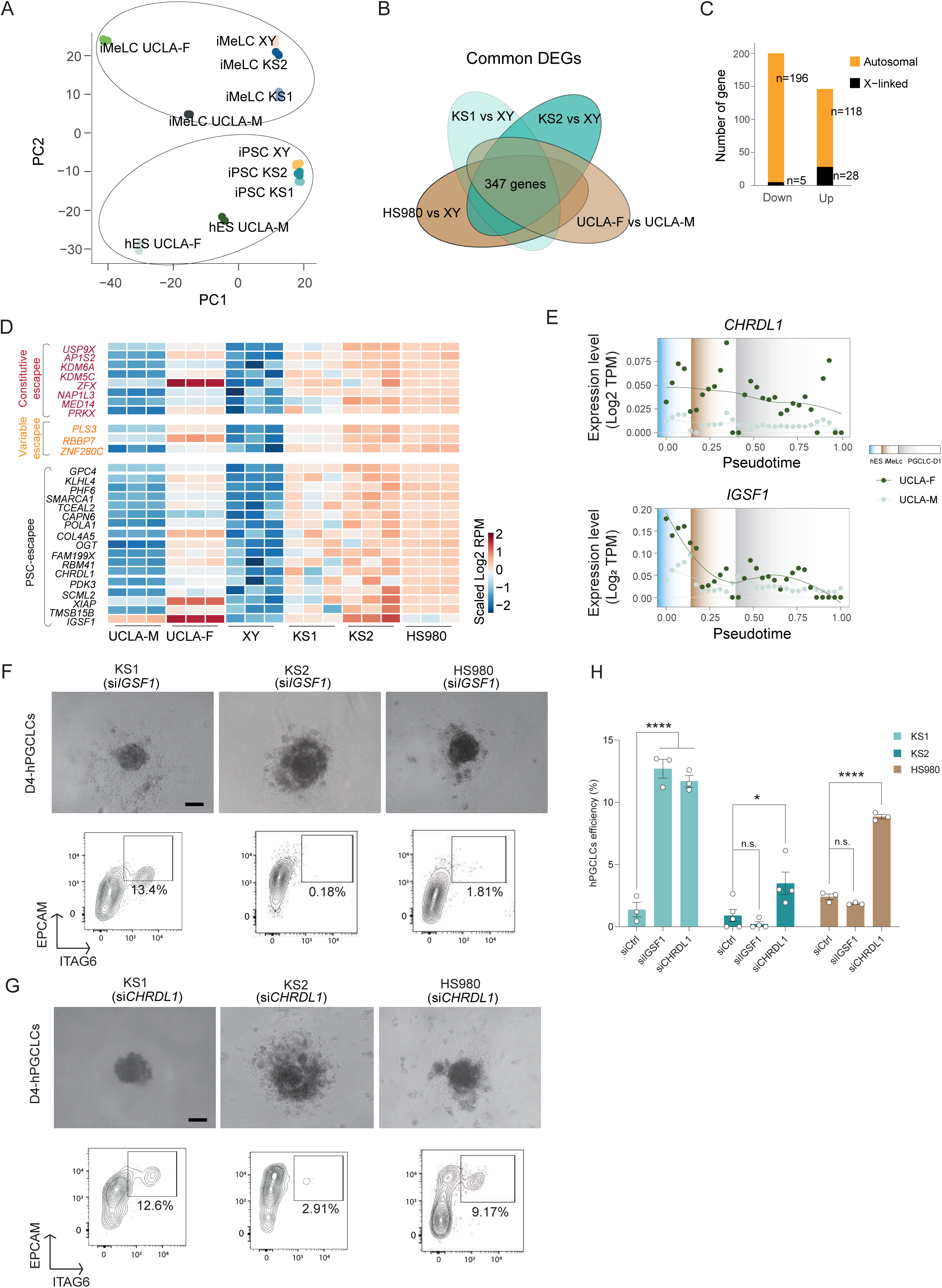
Downregulation of escape genes increased hPGCLC induction efficiency. (A) PCA plot showing similarities of KS1, KS2, XY, H9, HS980, UCLA-F, and UCLA-M at the PSC and iMeLC stages. (B) Venn diagram showing commonly dysregulated genes of the distinct KS or female lines vs XY line. (C) Gene count of the up- and down-regulated DEGs on the X chromosome and autosomes. (D) Heatmap showing gene expressions of PSC-related escapees. (E) Pseudotime associated gene expressions of IGSF1 and CHRDL1 from hES to iMeLC and D1 PGCLC in UCLA lines, respectively. TPM means Transcripts per million (F) D4 hPGCLC spheroid and FACS identification of hPGCLC after *IGSF1* KD. (G) D4 hPGCLC spheroid and FACS identification of hPGCLC after *CHRDL1* KD. (H) Statistics of D4 hPGCLC efficiencies after IGSF1 and CHRDL1 KD, respectively. (H) Pseudotime associated gene expressions of IGSF1 and CHRDL1 from hES to iMeLC and D1 PGCLC in UCLA lines, respectively. Data in (H) are shown as mean ± SEM. n.s., non-significant; \**p* < 0.05; \*\*\*\**p* < 0.0001 by one-way ANOVA comparison.

Next, we analysed differentially expressed genes (DEGs) at the PSC stage between each KS line and the control XY line, as well as between male and female lines, i.e., HS980 to XY, and UCLA-F to UCLA-M. As depicted in the Venn diagram, 347 DEGs were shared among four comparisons (Figure 2B, Supplementary Table 1), with 37 DEGs located on the X-chromosome and 314 DEGs on autosomes (Figure 2C). The X chromosome has a 5.6-foldchange in upregulated versus downregulated DEG counts, while autosomes ranged from 0.23 to 1.4, confirming the elevated X-linked dosage in the KS and female lines. Notably, three out of the 37 mitochondrial DNA encoded genes were commonly downregulated (Supplementary Table 1). Among the X-linked DEGs, 28 genes were up-regulated, including eight previously identified as constitutive escapees, such as *USP9X* and *KDM6A*,^5^ and three as variable escapees, such as *PLS3*, *RBBP7*, *ZNF280C* (Figure 2D).^5^ The remaining 17 genes were previously uncategorized and are herein considered as PSC-related escapees (Figure 2D). Within the escapees, we noticed that two genes, *CHRDL1* and *IGSF1*, could potentially be related to PGC induction. *CHRDL1* encodes an antagonist of BMP4^22^, while *IGSF1* is a negative modulator of the TGFβ-Activin pathway.^25^ Both BMP4 and TGFβ-Activin pathways play crucial roles in germ cell development and hPGCLCs induction. We further compared temporal gene expressions of CHRDL1 and IGSF1 during hPGCLC induction between male and female using pseudotime trajectory analysis. Utilizing previously available single-cell RNAseq data from iMeLCs and day1 (D1)-hPGCLCs on UCLA lines,^20^ our analyses revealed that the expression difference between UCLA-F and UCLA-M was observed both for *CHRDL1* and *IGSF1*, but the difference gradually diminished in iMeLCs and D1-hPGCLCs as the expression of both genes decreased (Figure 2E).

Subsequently, we examined whether knocking down (KD) each of these two escape genes could enhance the efficiency of hPGCLCs induction. Using siRNAs, we achieved ∼90% and ∼98% reduction of *IGSF1* and *CHRDL1* expression, respectively (Figure S3A). We then applied this KD approach in KS1, KS2 and HS980 during hPGCLCs differentiation. The *IGSF1* KD in the KS1 line resulted in an increased hPGCLC efficiency to around 13% (Figures 2F and 2H). While this efficiency is comparable to that of the normal female H9 line, it remained considerably lower than the XY control. Moreover, *IGSF1* KD did not significantly improve the efficiencies in KS2 and HS980 lines (Figures 2F and 2H). In contrast, *CHRDL1* KD not only significantly increased the efficiency in KS1 similar to *IGSF1*, but also greatly enhanced the efficiency in both KS2 and HS980 (Figure 2G and 2H). Noteworthily, despite the improvements after KD, none of the KS cell lines reached the efficiency levels comparable to that of the XY line. This suggests that other underlying mechanisms remain to be identified and understood.

### Upregulation of SOX2 by escapee gene *USP9X* hampers the hPGCLC induction

Interestingly, in addition to the X-linked genes, *SOX2* was among those upregulated autosomal DEGs (Supplementary Table 1). *SOX2* is well-known to be not conserved in human and mouse germ cell specification, but so far, the underlying mechanisms have remained elusive. We first corroborated that gene expression of *SOX2* increased in KS lines, HS980 and UCLA-F (Figure 3A), aligning with the increased SOX2 protein levels in KS lines at both iPSC and iMeLC stages compared to the XY line using immunofluorescence (Figures 3B and 3C) and Western blotting (Figure 3D). Importantly, SOX2 protein levels displayed failure to downregulation from iPSCs into iMeLCs, especially in the KS and HS980 lines compared to the XY and H9 line (Figures 3B and 3D). This elevated SOX2 level could pose a potential challenge for PGC induction, considering the documented reduction of SOX2 preceding PGC emergence. ^24, 26^ However, to date, concrete evidence directly linking *SOX2* with the inhibition of human germline specification is lacking.

**Figure 3.**
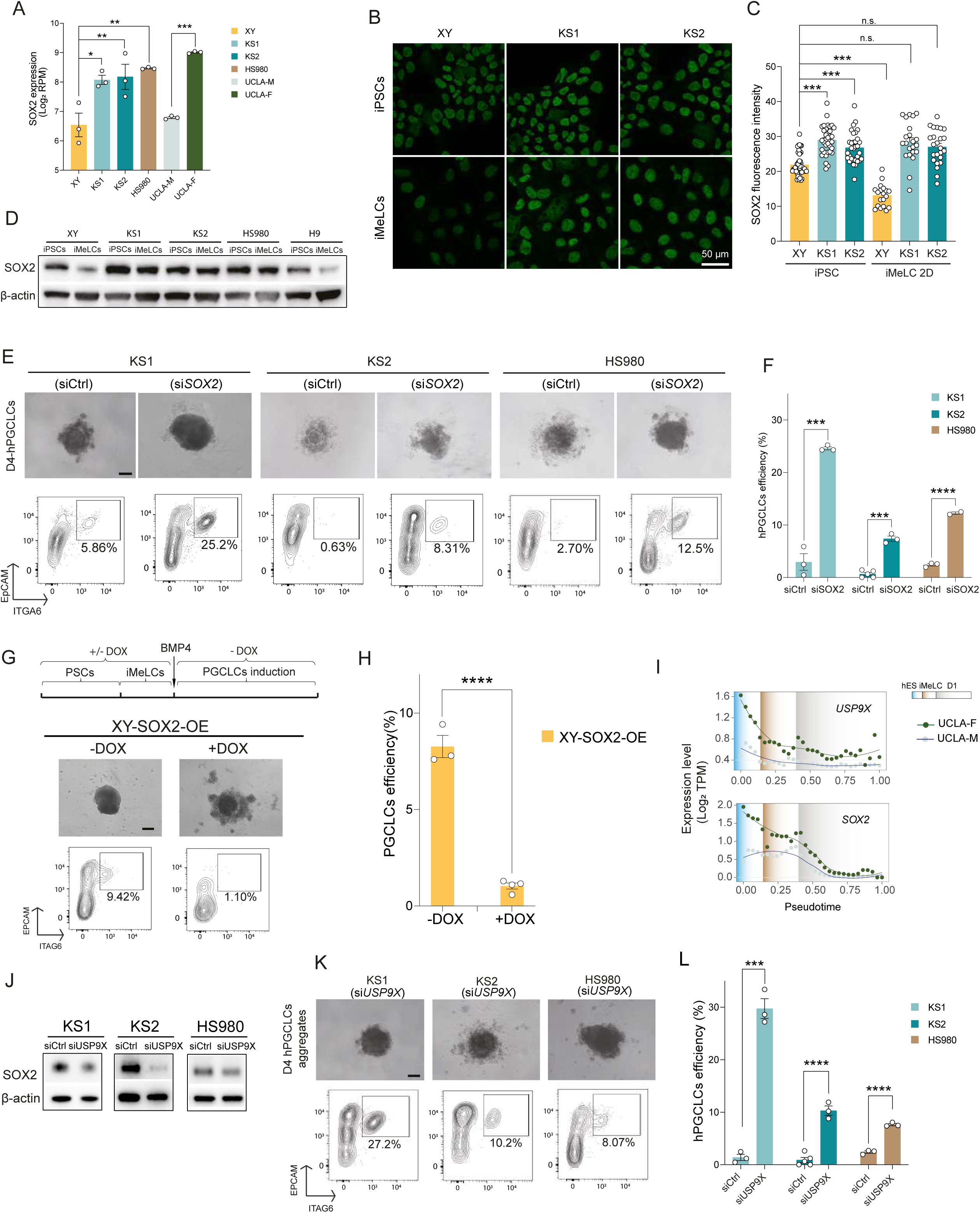
Upregulation of SOX2 by escapee gene *USP9X* interferes with the hPGCLC induction. (A) Normalized SOX2 gene expressions in the PSCs of XY, KS1, KS2, HS980, UCLA-F and UCLA-M. TPM means Transcripts per million. (B) Immunostaining showing SOX2 protein levels in the PSCs of XY, KS1 and KS2. Scale bar, 50 µm. (C) Fluorescence statistics analysis of SOX2 immunostaining. (D) Western blot showing SOX2 protein levels in the PSCs and iMeLCs of XY, KS1, KS2, HS980 and H9. (E) D4 hPGCLC spheroid and FACS identification of hPGCLC after SOX2 KD in KS1, KS2 and HS980, respectively. Scale bar, 200 μm. (F) Statistics of D4 hPGCLC efficiencies after SOX2 KD in KS1, KS2 and HS980 from three independent experiments. (G) Schematic diagram of DOX induces SOX2 OE, D4 hPGCLC spheroid, and FACS identification of hPGCLC after SOX2 OE in XY cells. Scale bar, 200 μm. (H) Statistics of D4 hPGCLC efficiencies after SOX2 OE in XY cells. (I) Temporal gene expressions of SOX2 and USP9X in UCLA-F and UCLA-M during hPGCLC induction. (J) Western blot showing SOX2 protein levels after USP9X KD. (K) D4 hPGCLC spheroid morphology and FACS identification of hPGCLC after *USP9X* KD in two KS and HS980 lines. Scale bar, 200 µm. (L) Summary of D4 hPGCLCs efficiency after USP9X KD in two KS and HS980 lines. Data in (A) and (C) are shown as mean ± SEM. \**p* < 0.05; \*\**p* < 0.01; \*\*\**p* < 0.001 by one-way ANOVA comparison. Data in (F), (H) and (L) are shown as mean ± SEM. \**p* < 0.05; \*\*\**p* < 0.001; \*\*\*\**p* < 0.0001 by t test.

To understand this, we first conducted *SOX2* KD using siRNA at the PSC stage in both the two KS lines and the HS980 line. Given that *SOX2* plays a crucial role in the pluripotency network in PSCs, our goal was to achieve a modest reduction in *SOX2* levels comparable to those found in male cell lines. Following optimization, we identified a dosage that consistently resulted in approximately 50% reduction of *SOX2* expression by Western blotting at the iMeLC stage (Figures S3B and S3C). We also monitored the colony morphology 72h after *SOX2* KD to ensure that the moderate *SOX2* KD did not disrupt their pluripotency maintenance (Figure S3D). Subsequently, we assessed the hPGCLC induction efficiency. In both the KS and HS980 lines, the yield of D4-hPGCLCs increased approximately fivefold in the *SOX2* KD groups compared to their control lines (Figures 3E and 3F). Notably, KS1 exhibited a yield of approximately 25%, comparable to the XY line (Figures 3E left panel and 3F). HS980 produced a yield of approximately 12.5%, comparable to the female control H9 line (Figures 3E and 3F). Although the yield of hPGCLCs of KS2 remained lowest, it was comparable with other previously reported female lines (Figures 3E and 3F).^19, 20^

Next, we examined whether SOX2 overexpression (OE) in the control XY line would correspondingly reduce hPGCLC differentiation efficiency. To accomplish this, we established an inducible XY-SOX2-OE line through lentivirus transduction and puromycin selection, in which SOX2-OE could be induced by the doxycycline (DOX) administration. As a result, we observed failure of spheroid aggregation in responsive to elevated SOX2 levels in the +DOX group after BMP induction (Figures 3G). Consequently, the capacity for PGCLCs differentiation was also lost in the +DOX group compared to the -DOX group (Figure 3G and 3H). These observations were reminiscent of the spheroid morphology and hPGCLCs formation found in the KS and HS980 lines (Figure 1B and 1C).

As the SOX2 elevation caused reduced PGCLC induction in the KS and female lines, we investigated whether the *SOX2* upregulation in the KS and female lines is associated with XCI escape genes. Previous studies have identified that *USP9X* upregulates SOX2 at both mRNA and protein levels.^27^ Moreover, *USP9X* is the most abundantly expressed escape gene that was located in a region showing the most gene expression on the X chromosome (Figures 1G and 2D). Furthermore, using pseudotime analysis revealed a similar pattern of gene expression for *USP9X* and *SOX2*, including their rapid reduction during the differentiation process and the male-to-female differences (Figure 3I). Therefore, we examined whether *USP9X* KD would lead to decreased *SOX2* levels. Indeed, *USP9X* KD reduced *SOX2* expression at mRNA level in iMeLCs by approximately 20%, 40% and 30% in KS1, KS2 and HS980, respectively (Figure S3E). USP9X KD also decreased SOX2 protein levels at the iMeLC stage, confirmed by Western blotting (Figure 3J). Notably, like *SOX2* KD, *USP9X* KD resulted in higher hPGCLC yields compared to previous *IGSF1* KD and *CHRDL1* KD (Figure 3K). The *USP9X* KD not only increased the efficiency of KS1 to the level of the control XY line but also enhanced the efficiency of KS2 and HS980 to levels similar to H9 line (Figures 3K and 3L). The proliferation activities of hPGCLCs typically peak at D4, with hPGCLCs efficiency greatly declines at D6.^24^ Interestingly, both SOX2 KD and US9X KD led to a sustained high yield of hPGCLCs at D6 (Figures S3F-S3H), highlighting the regulatory axis of USP9X-SOX2 against the germline induction.

### SOX2 interferes with mitochondrial oxidative metabolism

Interestingly, in addition to SOX2, we observed that many commonly downregulated DEGs in KS were autosomal genes (Figure 2B), related to mitochondrial function (total 55 genes), oxidative phosphorylation and aerobic respiration (total 80 genes), including mitochondrial cytochrome c oxidase (COX) genes and NADH oxidoreductase core subunits (NDUF) genes (Figure 4A, Figure S4). Notably, genes associated with mitochondrial morphology dynamics, such as OPA1 and GDAP1, were upregulated. Among mitochondria related DEGs, 35 were downregulated and 20 were upregulated in the KS lines compared to the XY control. For aerobic respiration related DEGs, the majority (69 out of total 80 genes) were downregulated in the KS lines (Figure 4B), indicating an enrichment of reduced function. Gene set enrichment analysis (GSEA) further confirmed that pathways related to aerobic respiration were significantly downregulated in the KS and female lines, including ATP synthesis and its coupled electron transport, and oxidative phosphorylation (Figure 4C, Supplementary Table 2).

**Figure 4.**
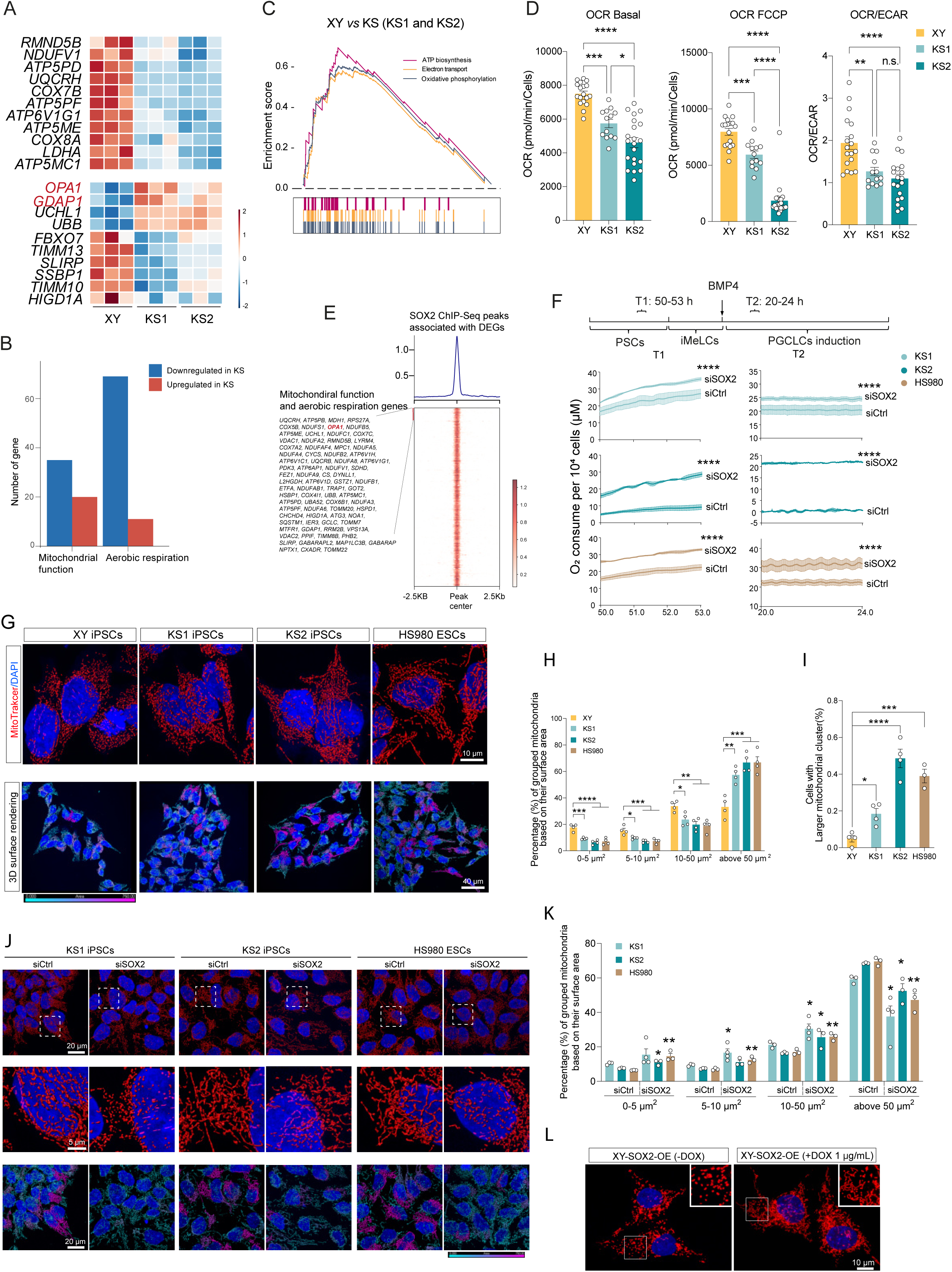
SOX2 interferes with cellular bioenergy metabolism and mitochondrial morphology dynamics. (A) Heatmap showing selected dysregulated genes related with oxidative phosphorylation and mitochondria in KS compared to XY. (B) Gene counts of commonly up- and down-regulated genes related with oxidative phosphorylation and mitochondria in KS compared to XY. (C) GSEA result showing decrease of oxidative phosphorylation related pathway in KS. (D) Seahorse assay showing that KS1 and KS2 had lower oxidative phosphorylation than XY. (E) Heatmap showing the SOX2 ChIP-seq peaks associated with the DEGS of the two KS lines vs the XY line. (F) Resipher assay showing the changes of O_2_ consumption after SOX2 KD in selected time window in the KS and HS980 lines, in their PSC stage (T1) and during their hPGCLC induction (T2), respectively. (G) Mitochondrial morphology via mitoTracker staining and 3D rendering in the PSCs of XY, KS1, KS2 and HS980 lines. (H) Categorized mitochondrial surface area from three independent experiments. XY, n=1731; KS1, n=2429; KS2, n=993; H9, n=1386; HS980, n=1782. N denotes to number of mitochondria used for the surface area estimation. (I) Percentage of cells with mitochondria clusters. Mitochondria cluster is defined as surface area larger than 500 µm^2^. (J) Mitochondria morphology via mitoTracker staining and 3D rendering in the PSCs of KS1, KS2 and HS980 after SOX2 KD. (K) Percentages of cells with mitochondria clusters after SOX2 KD from three independent experiments. KS1 siCtrl, n=2697; KS1 siSOX2, n=2261; KS2 siCtrl, n=2014; KS1 siSOX2, n=5687; HS980 siCtrl, n=5312; HS980 siSOX2, n=9015. N denotes to number of mitochondria used for mitochondria cluster estimation. (L) Mitochondrial morphology via mitoTracker staining in the PSCs of XY-SOX2-OE before and after SOX2 OE. Data in (D), (H), (I), (K) are shown as mean ± SEM. n.s., non-significant; \**p* < 0.05; \*\**p* < 0.01; \*\*\**p* < 0.001, \*\*\*\**p* < 0.0001 by one-way ANOVA comparison. Data in (F) are shown as mean ± SEM. \*\*\*\**p* < 0.0001 by two-way ANOVA comparison.

To further examine the functional consequences of the downregulation of these mitochondrial genes, we performed Seahorse XF Cell Mito Stress tests on XY, KS1, KS2 and HS980 lines. After normalizing the cell population sizes, we found that KS and HS980 PSCs displayed low oxygen consumption rates (OCR) under the basal and stressed conditions, indicating impaired oxidative phosphorylation (Figures 4D and S5A-S5C). In addition, the KS lines showed a shift toward glycolysis, as evidenced by a lower ratio of OCR to extracellular acidification rates (ECAR) compared to the XY control (Figure 4D). These findings corroborated with a lower proliferation rate in KS lines, especially KS2 (Figure S5D).

SOX2 has been shown to mediate metabolic reprogramming in cancer cells by enhancing glycolysis.^28–30^ To examined whether elevated SOX2 accounted for dysregulation of mitochondrial function, we first analyzed SOX2 binding sites using published ChIP-seq data from human iPSCs^31^ and obtained 3741 peaks after filtering out peaks with enrichment scores under 25. These peaks were annotated to transcription starting sites (TSS) of genes within 125 kb interval, the reported median distance between enhancers and promoters in the human genome.^32^ Comparing these annotated genes with the commonly dysregulated 2238 DEGs in KS1 and KS2 lines, we identified 1328 DEGs potentially regulated by SOX2 (Figure 4E).

Among these genes, 79 genes were associated with mitochondrial function, including OPA1 and oxidative phosphorylation related genes (Figure 4E). To confirm the causal association between SOX2 and oxidative phosphorylation, we used Resipher device to measure oxygen (O_2_) consumption in real time in KS1, KS2 and HS980. We compared O_2_ consumption per 10^4^ cells in *SOX2* KD with control siRNA during two critical periods: T1 (50-53h after PSC culture passage) and T2 (20-24h after hPGCLCs induction). We found that O_2_ consumption was significantly increased in all three lines upon SOX2 KD (Figure 4F). Among them, KS2 exhibited the lowest O_2_ consumption, especially during T2 period, consistent with the failure of spheroid formation and extensive cell death observed previously (Figures 1B and S1C). Importantly, *SOX2* KD resulted in a significant increase in O_2_ consumption from nearly zero to approximately 21 µM per 10^4^ cells (Figure 4F).

### SOX2 interferes with mitochondria morphology

Previous studies have shown that stem cell maintenance and differentiation are intrinsically linked with energy metabolism.^33–35^ Pluripotency culture conditions intrinsically program bioenergetic metabolism, which in turn regulates the epigenetic mechanisms to consolidate the pluripotency state and differentiation potential.^36, 37^ In response to cellular metabolism demand, mitochondria undergo morphology transitions, regulated by mitochondrial fusion and fission dynamics.^33, 38^ We examined mitochondrial dynamics related genes from our sequenced data and found *OPA1*, which promotes mitochondrial fusion, significantly upregulated in the two KS lines (Figure 4A and Figure S5E). In addition, the elevated expression of OPA1 is likely related to SOX2 binding (Figure 4E). We therefore examined the mitochondrial morphology using mitotracker staining. Mitochondria in PSCs often display rounded and fragmented morphology^39^ and in agreement with this, we found that mitochondria appeared most fragmented and spherical in XY cells compared with other lines (Figure 4G). In contrast, mitochondria in two KS lines, especially KS2, and HS980 line displayed more fused and tubular appearance at both PSC and iMeLC stages (Figures 4G and S5F). Also, compared to KS1, a greater number of cells in KS2 and HS980 exhibited increased mitochondrial clustering, distinct from control XY and H9 lines at both PSC and iMeLCs stage (Figures 4G and S5G). These observations were further strengthened by analysis of 3D surface rendering of confocal images using Imaris software, where larger mitochondrial clusters were represented in pink (Figure 4G lower panel and S5G).

To further analyze mitochondrial dynamics, we utilized Imaris software to quantify the size of individual mitochondria and mitochondrial clusters in each cell line based on their surface area. We categorized mitochondria into four groups depending on their surface area (Figure 4H). Our results showed that both KS lines and HS980 exhibited a significantly decreased proportion of the smallest mitochondria (0-5 µm^2^), with KS2 having the lowest number (Figure 4H). Notably, both KS2 and HS980 also contained a lower number of middle-sized mitochondria (5-10 µm^2^ and10-50 µm^2^) (Figure 4H). Additionally, KS2 and HS980 showed a significant increase in the size of larger mitochondrial clusters (>50 µm^2^). Moreover, KS2 and HS980 cells also displayed a significantly higher degree of mitochondrial clustering, defined as the ratio of larger mitochondrial clusters (area > 500 µm^2^) relative to total mitochondrial surface area (Figure 4I). Specifically, the percentage of cells with larger mitochondrial clusters (indicated in pink in Figure 4G) was significantly higher in KS1 (19.3 ±2.92), particularly in KS2 (46.3 ±5.04) and HS980 (38.9 ±5.25) compared to XY (5.40 ± 2.46) (Figure 4I). This severe mitochondrial clustering persisted into the iMeLCs stage (Figure S5G and S5H).

Considering that SOX2 KD resulted in increased oxygen consumption, we investigated whether *SOX2* KD could regulate mitochondrial fission and fusion dynamics. To confirm this, we assessed the mitochondrial morphology using Mitotracker staining following *SOX2* KD. We observed a transition from fused and clustered mitochondria to more rounded and fragmented mitochondrial shapes in the KS1, KS2 and HS980 cell lines treated after S*OX2* KD, compared to controls (Figure 4J, upper and middle panels, and 4K). Additionally, there was a substantial decrease of mitochondrial clustering especially in KS2 and HS980 (Figures 4J, lower panel and 4L). Specifically, the percentage of cells with larger clustered mitochondria significantly decreased from an average of 20% in the control group to 4% in the KS1 SOX2 KD. Remarkably, it dramatically decreased from 50% in both to 15% and 10% in KS2 and HS980, respectively (Figure 4L). In contrast, overexpressing SOX2 in XY led to fused and elongated mitochondria (Figure S5I).

To further understand the potential molecular regulation of mitochondrial morphological dynamics, we plotted the key genes involved in regulating mitochondrial fission, fusion, mitophagy and mitochondria trafficking. Among these, we found consistent upregulation of the fission gene *GDAP1*, the fusion genes *OPA1* and *YME1L1*, and the mitochondria trafficking genes *KIF5A*, *KIF5AC* and *RANBP2* in both KS and HS980 (Figure S4I).

### Timely SOX2 silencing is critical for human germline fate determination

PGCLC induction coincides with gastrulation, during which progenitors transiently acquire a mesoendoderm-like identity. Previous studies have shown that increased SOX2 and glycolysis have been shown to favor ectoderm/neuroectoderm lineage specification.^40–43^ Therefore, we asked whether iMeLCs with elevated SOX2 and reduced oxidative phosphorylation were predisposed towards the ectoderm lineage. To this end, we performed Smartseq3 single-cell RNA-seq on XY, KS1 and K2 at the iMeLC stage. After plotting these cells together with UCLA-F and UCLA-M into the PCA, we observed the primary segregation was between all male lines (XY, KS1, KS2, UCLA-M) and female UCLA-F line (Figure 5A). To evaluate the cell fate transition, we applied Capybara quadratic programming, a recently developed tool to measure cell identity at single-cell resolution to evaluate the cell fate transition.^44^ Our analysis revealed a marked increase in ectoderm identity and a decrease in mesoendoderm identity in UCLA-F and the two KS lines compared to male controls (Figure 5A). These differences in cell identity among all lines were further confirmed by the cell density plot (Figure 5B). Using the same analysis, we found that *SOX2* KD notably shifted cell fates towards the mesoendoderm identity in both two KS and HS980 cell lines compared to controls (Figure 5C). These findings suggested that *SOX2* acts as an inhibitor of mesoendodermal fate, consequently hampering hPGCLCs induction.

**Figure 5.**
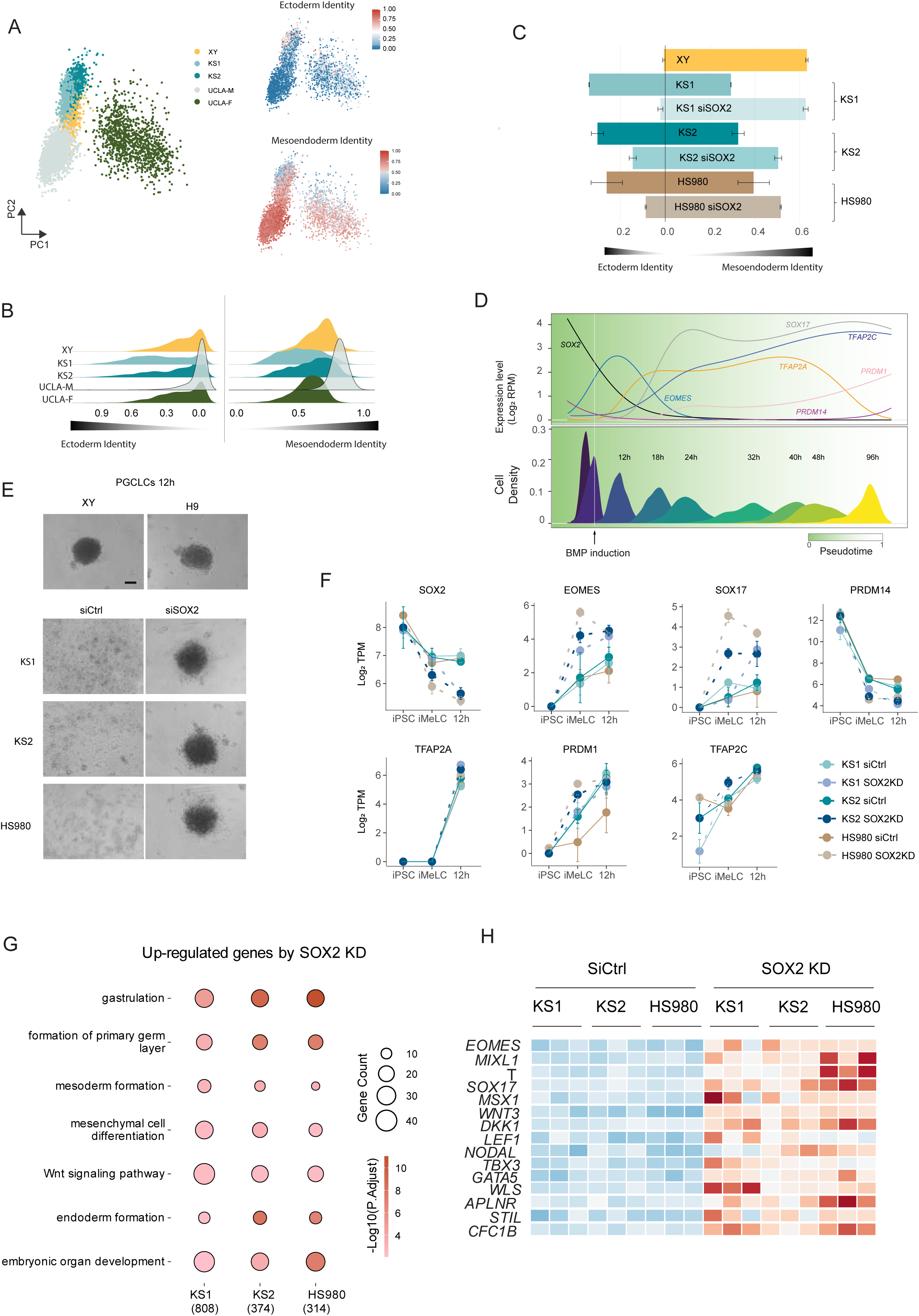
SOX2 predisposes cells towards the ectoderm fate. (A) Ectoderm and mesoendoderm identities of the iMeLCs of XY, KS1, KS2, UCLA-M and UCLA-F. Left, PCA of the iMeLCs of XY, KS1, KS2, UCLA-M and UCLA-F. Right upper, Ectoderm identities in the same PCA embedding. Right lower, mesoendoderm identities in the iMeLCs of each cell line. (B) Density plot of the ectoderm and mesoendoderm identities in the iMeLCs of each cell line in A. (C) Ectoderm and mesoendoderm identities changes in the iMeLCs of KS1, KS2 and HS980 after SOX2 KD. (D) Temporal gene expressions of SOX2, SOX17, TFAP2A, TFAP2C and RPDM1 during male hPGCLC induction. TPM means Transcripts per million. (E) Represent morphology of PGCLCs 12h spheroid of XY, H9, KS1, KS2 and HS980 before and after SOX2 KD. Scale bar, 200 µm. (F) Selected gene expression changes after SOX2 KD in the iMeLC and 12h post BMP induction stages in KS1, KS2 and HS980. TPM means Transcripts per million. (G) Common up regulated pathways after SOX2 KD in the 12h stage of KS1, KS2 and HS980. KS1, KS2 and HS980 has 808, 374 and 314 DEGs after SOX2 KD, respectively. (H) Heatmap showing the changes of selected genes after SOX2 KD in the 12h stage of KS1, KS2 and HS980.

Upon hPGCLC induction, BMP signaling activates TFAP2C in a SOX17-independent manner meanwhile represses SOX2.^24, 26^ We confirmed this by pseudotime analysis of UCLA-F and UCLA-M where SOX2 diminished at the end of D1 after BMP treatment in both lines (Figure 3I). Thus, we investigated whether elevated SOX2 in the KS and HS980 impacted the timing of SOX2 silencing post-BMP induction, specially comparing the duration from BMP addition to D1 comparing XY control with KS and HS980 lines.To this end, we examined the temporal expression of *SOX2* during hPGCLCs induction. We analyzed a recently published single-cell dataset containing several time points, including 12h, 18h and 24h to D4 after hPGCLC induction.^45^ We constructed the pseudotime differentiation trajectory across the entire differentiation process and charted the *SOX2* expression together alongside several key genes for the germline specification, i.e., *SOX17*, *TFAP2C*, *TFAP2A* and *PRDM1.* We found that SOX2 begins to decrease during the iMeLC stage and becomes completely repressed at 12 h after BMP4 induction, preceding the emergence of the key regulators *TFAP2A* and *SOX17* for the hPGCLC lineage (Figure 5D). Furthermore, the expression of *TFAP2C* and *PRDM1* gradually increased after 24h (Figure 5D). These temporal gene expressions suggested that 12h after BMP induction is a critical point for SOX2 repression. Persistently high levels of *SOX2* expression could interfere with the germline fate induction by hindering the initiation of these key genes after BMP4 induction.

Therefore, we examined SOX2 expression at this time point. Unlike the XY line, where SOX2 was essentially silent at D1, the KS lines exhibited a 2.6-fold higher expression of SOX2 at 12h after BMP induction (Figure S6A). Additionally, the KS and HS980 lines failed to form spheroids at 12h, in contrast to the XY and H9 lines (Figure 5E), which was notably improved by SOX2 KD (Figure 5E). To unravel the mechanism, we analyzed the expression of SOX2 and other lineage determinant markers. We found that *SOX2* KD slightly reduced its own expression at the iMeLC stage and silenced it at 12h, in contrast to the control group (Figure 5F). Interestingly, the mesoderm marker *EOMES* and the PGC marker *SOX17* were drastically increased after SOX2 KD (Figure 5F). Additionally, PRDM14 was decreased by *SOX2* KD at 12h (Figure 5F), corroborating previous findings that *PRDM14* needs to be repressed following BMP treatment for PGC induction during the first 2 days, and reactivated once PGCs establish their pluripotency network.^26^ Other important TFs, including *TFAP2A*, *PRDM1*, *TFAP2C*, did not show differences (Figure 5F).

Given that 12h is the critical time point for hPGCLC induction, we compared the enriched upregulated pathways after SOX2 KD and found consistent enrichment in pathways related to gastrulation, mesoderm and endoderm development across all cell lines (Figure 5G). Specifically, the genes upregulated by SOX2 KD included key genes such as *NODAL*, *LEF1*, *WNT5B*, *WNT3*, *TBX3*, *GATA5*, *FGF17*, *SOX17*, as well as mesoderm progenitor genes *EOMES*, *MIXL1*, and *T* (Figure 5I, Supplementary Table 3). Downregulated genes included *SOX2* itself, genes in the NODAL and WNT pathway such as *LEFTY2* and *TCF7L1* (Supplementary Table 3). Interestingly, the KS1 line exhibited an increase in multiple pathways related to oxidative phosphorylation 12h after SOX2 KD during hPGCLC induction (Figure S6B).

### Elimination of SOX2 is a prerequisite for SOX17 expression

The molecular cascades and signalling pathways involved in germ cell fate determination are generally conserved between mice and humans, with a notable divergence in term of the role of SOX17 in human instead of SOX2 in mice.^26^ The mechanistic implications of this divergence are not well understood, with speculation focusing on difference in embryonic structure and early pluripotency states. Our SOX2 KD experiment demonstrated that a reduction in SOX2 levels significantly elevated SOX17 (Figure 5F). We therefore speculated that the repression of SOX2 is a prerequisite for the expression of SOX17 in humans. To this end, we assessed SOX2 and SOX17 protein levels using FACS in D1-PGCLCs. In the XY control, we found about 40% SOX17-positive cells and around 2% SOX2-positive cells, with no double positive cells detected (Figure 6A). In contrast, the two KS and HS980 lines exhibited higher residual SOX2-positive cells (around 5%) and lower SOX17-positive cells (around 20%), also with no double positive cells are present (Figure 6A and S6C). This confirms that SOX2 is normally repressed before the onset of SOX17 and the delayed repression of SOX2 in the KS and female lines resulted in fewer SOX17 positive PGCLC progenitor cells.

**Figure 6.**
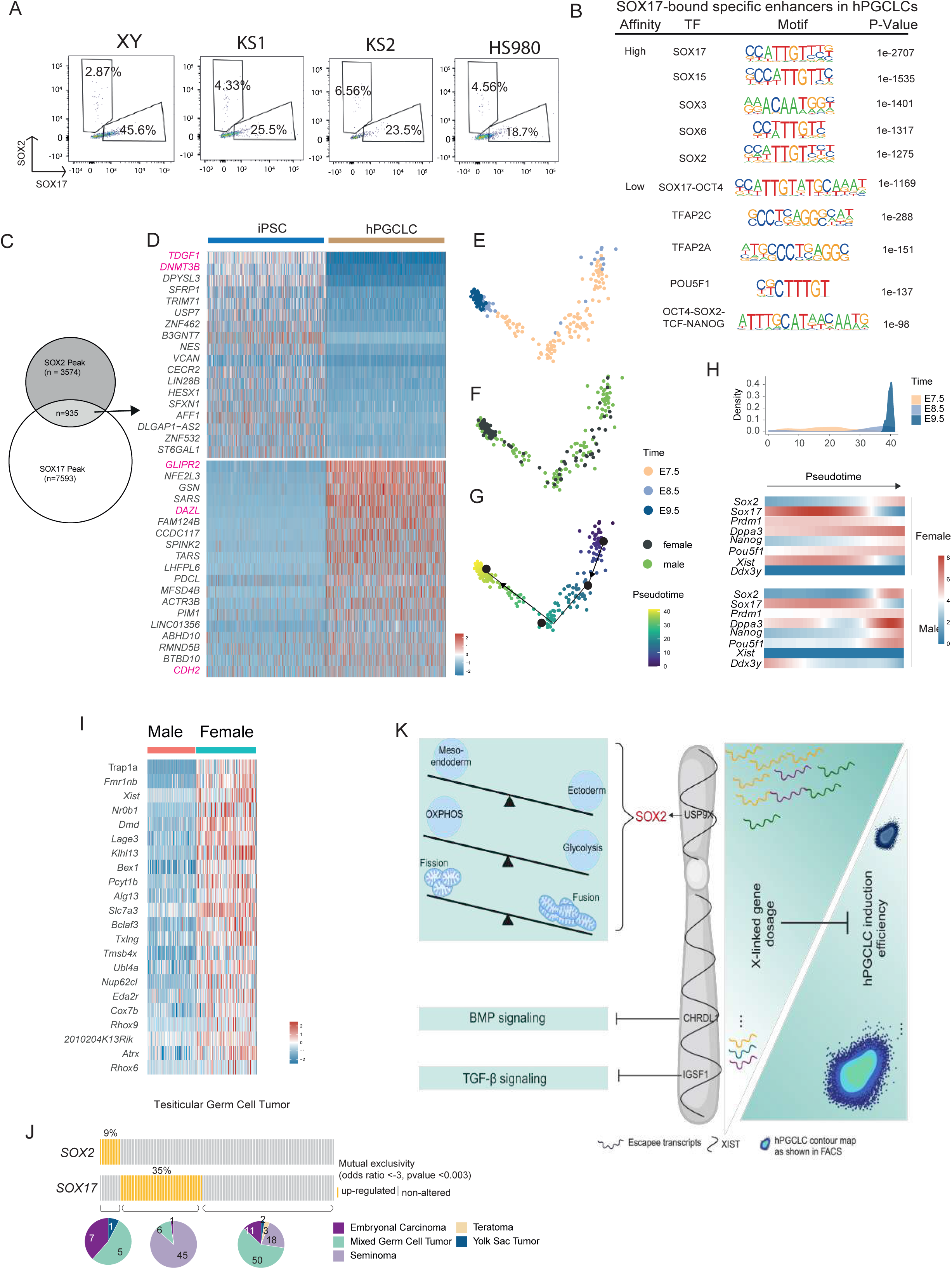
Elimination of SOX2 is a prerequisite for SOX17 expression. (A) FACS plot analysis of SOX17- and SOX2-expressing cells in the XY Day 1 hPGCLC spheroid. (B) Motif enrichments of the potential high-affinity and low-affinity SOX17 binding sites. (C) Venn diagram showing overlapping of SOX2 peaks in iPSC and SOX17 peaks in hPGCLCs. (D) Heatmap showing the genes that are differently regulated by SOX2 and SOX17 binding, respectively. (E-G) PCA plots showing E7.5-E9.5 mPGCs, with the information of developmental time points, sex and inferred pseudotime(H) Exclusive expression of Sox2 and Sox17 duringE7.5 to E9.5 mPGC progression. (I) X-linked genes upregulated in mouse female EpiSCs compared to male EpiSCs. (J) mutual exclusivity of SOX2 and SOX17 in human Testicular germ cell tumor, with the expression pattern and tumor type shown. (K) Schematic summary of how does increased X–linked gene dosages inhibit hPGCLC differentiation.

Next, we analysed the cis-regulatory elements (CRE) bound by SOX17 in hPGCLCs using a published ChIP-seq dataset.^46^ Using log-odds scores to evaluate the match between the peak sequences and the canonical SOX17 motifs, the Homer algorithm^47^ distinguished SOX17-bound CREs into high-affinity and low-affinity binding sites based on conservation information. We then conducted motif enrichment on these two groups of peaks. Aligning with their high-affinity characteristics, these peaks were highly enriched for SOX family motifs, with SOX17 being the most prominent (Figure 6B). Among the top five enriched motifs, we also identified the canonical motif for SOX2 (Figure 6B). Notably, only SOX2 but not SOX15, SOX3 and SOX6 was among our previously identified DEGs (Supplementary Table 1), suggesting that SOX2 but not the other SOX proteins could compete with SOX17 for binding downstream genes in hPGCLCs induction. The SOX17-low-affinity binding sites were enriched with motifs of SOX17-OCT4 cooperative motifs, TFAP2C and TFAP2A, further supporting our findings. Among the top enriched motif for low affinity binding, we also identified OCT4-SOX2-TCF-NANOG enrichment, indicating that SOX2 could also compete with SOX17 for low affinity binding sites (Figure 6B). These findings suggested that ectopic SOX2 expression might lead to the unintended occupation of CREs by SOX2 or SOX2-containing complexes, potentially excluding SOX17 from those binding sites and thereby altered downstream regulatory effects.

To further confirm the perturbation by SOX2 on SOX17 regulatory sites, we overlapped SOX2 binding sites in hiPSCs with SOX17 binding sites in hPGCLCs, and identified 935 common sites (Figure 6C). We then annotated these sites to genes and examined their expressions in iPSCs and hPGCLCs, focusing on genes with dynamic patterns, i.e., transitioning from no/low expression to high expression or vice versa. This approach could reflect opposing roles of of SOX2 and SOX17 in gene regulation. We identified 18 genes, such as *TDGF1* and *DNMT3B*, activated by SOX2 in iPSCs but silenced by SOX17 in hPGCLCs (Figure 6D). Conversely, we also found that 19 genes, including the EMT genes *GLIPR2* and *CDH2*, and the PGC gene *DAZL* were inhibited by SOX2 in iPSCs but activated in hPGCLCs (Figure 6D). This suggested that ectopic SOX2 expression may disrupt the proper silencing or activation of critical genes by SOX17, highlighting the necessity of SOX2 elimination for SOX17 to effectively regulate its downstream targets. A previous study from Yabuta et.al has identified a transient upregulation of *Sox17* around E7.25 in mouse PGCs.^48^ Recently, a high-precision transcriptome atlas of 11,598 mouse germ cells covering 28 critical developmental time points was generated.^49^ We therefore reanalyzed mouse PGCs from early developmental timepoints E7.5, E8.5 and E9.5 and performed clustering using PCA (Figure 6E). Noting that the majority of these PGCs presented exclusive gene expressions of *XIST* and Y chromosome gene *Ddx3y*, we further annotated these cells into 173 male and 93 female PGCs (Figure 6F and S6D), with their PGC identify further confirmed by double gene expression of *Prdm1* and *Dppa3* (Figure S6E).^48^ We then proceeded with the pesudotime analysis to align all PGCs along their developmental trajectory to examined *Sox2* and *Sox17* temporal expression (Figure 6G). When plotting the distribution of PGCs along the pseudotime trajectory, *Sox2* and *Sox17* were mutually exclusively expressed, with *Sox17* transiently expressed around E7.5 (Figure 6H). Interestingly, we found that female and male PGCs showed similar expression patterns and levels (Figure 6H). We also examined upregulated X-linked genes in female versus male mouse primed epiblast stem cells (mEpiSCs) when XCI is mostly completed. Unlike human iPSCs, only a few upregulated X-linked genes were identified (Figure 6I), among which *Usp9x* was not one of those in line with similar *Sox2* expression level in mouse PGCs (Figure 6I). Additionally, the other two human escapee genes *Igsf1* and *Chrdl1* were also not found. This lack of expression of these X-linked genes could underlie the specification programs of mouse PGCs with Sox2 versus human PGCs with SOX17.

To further investigate the mutual exclusivity of SOX2 and SOX17 in humans, we turned to human testicular germ cell tumors (TGCTs), the most common tumors in male aged 20-40 years.^50^ Among TGCTs, seminoma is the most prevalent type and shares gene expression programs with PGCs.^51, 52^ Unlike seminomas, nonseminomas encompass diverse tumor types, including embryonal carcinoma, yolk sac tumor, choriocarcinoma, teratoma.^51^ We analyzed the expression of SOX2 and SOX17 in 149 TGCTs samples with subtype information from The Cancer Genome Atlas Program (TCGA) and identified significant mutual exclusivity in terms of expression of SOX2 and SOX17 (Figure 6J). SOX2 expression is predominantly restricted to embryonic carcinoma, followed by mixed germ cell tumors and one case of yolk sac tumor, whereas SOX17 is mainly in seminoma with several cases of mixed germ cell tumors and one case of embryonic carcinoma (Figure 6J).

Taken together, SOX2 and SOX17 are mutually exclusive in human and mouse PGCs. Human escapee USP9X, which is not expressed from inactivated X chromosome in mice, maintains high SOX2 expression level during PGCLCs differentiation. SOX2 disrupts PGC fate establishment by occupying SOX17 functional positions and regulatory binding sites, competing for binding motif with SOX17 and suppressing genes activated by SOX17.

## Discussion

To achieve roughly equivalent gene expression outputs between female and male, epigenetically regulated silencing on one of the two X chromosomes occurs in female. Despite this, accumulating evidence indicates that XCI is frequently incomplete and X-linked gene dosage between male and female significantly affects complex trait variation.^5, 53, 54^ Our study has now unveiled the multifaceted effects of X-linked gene dosage on hPGCLCs specification, underlying not only the difference between male and female but also the disease mechanisms associated with supernumerary X chromosomes in KS. These effects are mediated not only by escape genes but also by autosomal genes, as X-linked genes can directly or indirectly regulate autosomal gene expression. Specifically, escape genes *IGSF1* and *CHRDL1* have been previously demonstrated to inhibit the TGF-beta/Activin A and BMP pathway, respectively, which are two essential signaling pathways required in PGC specification. More importantly, our findings revealed a connection between X-linked and autosomal genes, specifically *USP9X* regulating *SOX2* expression levels. Our findings suggested that *SOX2* expression can be influenced by human-specific escape genes, and elevated *SOX2* expression negatively impacts hPGCLCs specification. Our research underscores the importance of downregulting SOX2 expression during early human germ cell specification and provides insight into non-conserved function of SOX2 in PGC fate specification between mice and humans (Figure 6K).

### Human-specific X-linked genes affect PGCLC specification

Species-specific differences in the identity of escape genes have been implicated in the evolution of sex differences. In humans, about 15% of X-linked genes consistently escape XCI, whereas in mice, this proportion is only around 3%.^4, 55^ Given that sex differences in physiology and traits are generally more pronounced in humans than in mice, it is reasonable that a greater number of escape genes could provide an evolutionary advantage. Indeed, escape genes play important roles, as exampled by Turner syndrome, which manifests severe reproductive and neurological phenotypes as well as several physical abnormalities.^56^ Our results revealed that three escape genes are associated with hPGCLC specification. While female PSCs with proper XCI still generate hPGCLCs, the efficiency is compromised.^19, 20^ However, an increased dosage of X-link genes, due to erosion of XCI or supernumerary X chromosomes, could severely interfere with hPGCLC induction. We focused on *SOX2* because its function in PGC fate specification is not conserved between mice and humans. Also, *SOX2* has been reported to be directly or indirectly regulated by these three escape genes.^22, 25, 27, 57, 58^ Especially, *USP9X* encodes a ubiquitin-specific protease and plays an important regulatory role in protein turnover by preventing degradation of proteins through the removal of conjugated ubiquitin.^59^ USP9X has been well studied for its roles in neural developmental disorders and cancer.^60^ In osteosarcoma, USP9X serves as a bona fide deubiquitinase for SOX2, which is a critical oncoprotein in osteosarcoma.^61^ Similarly, USP9X KD in melanoma results in elevated ubiquitination of SOX2, subsequently suppressing tumor growth.^27^ Therefore, targeting the USP9X/SOX2 axis has been proposed as a novel strategy for SOX2-related cancers. Our study suggested that the USP9X/SOX2 axis is also involved in the specification of hPGCLCs, imposing an additional regulatory layer in females. A distinct genetic regulatory network in human PGC fate specification with SOX17 may have invented to counteract this challenge.

It would be of great interest to understand whether there are differences in the early human PGCs specification between female and male, but it is challenging to gain access to such early stage of human embryos. It is highly plausible that the current *in vitro* culture conditions do not fully replicate the optimal *in vivo* environment, where additional regulatory factors may compensate for X-linked gene dosage effects on human PGCs specification. Despite these limitations, our findings with *SOX2* KD open the possibility for generating hPGCLCs even from hPSCs with unfavourable XCI states.

### Sex and species difference in PGCs specification

It is generally considered that there are no transcriptional differences between male and female PGCs until sex determination initiates after gonadal entry.^62, 63^ However, PGC development in mice is more synchronized compared to that in humans. In humans, asynchronous and heterogeneous development is more pervasive in female fetal germ cells.^64^ Even at later developmental stages (e.g., week 26), early mitotic germ cells still co-exist with three other later stages of fetal germ cells.^64^ The relationship between this heterogeneous development in female germ cells and the epigenetic reprogramming of the X chromosome remains elusive. In mice, the dynamic of X chromosome activity and dosage compensation between male and female have been extensively studied.^65^ XCI occurs in epiblast cells before PGCs specification and is maintained during PGCs migration.^66^ Although one X chromosome is silenced in PGCs, the total gene output from X chromosomes is slightly higher in female mouse PGCs than in male mouse PGCs, which also appears to be conserved in humans.^65^ Interestingly, the Y chromosome can influence X dosage compensation states during PGC specification.^65^ Therefore, understanding the dynamics of the X chromosome activity during female germ cell development is important in both mice and humans. When establishing *in vitro* culture conditions, careful consideration must be given to the intricacies associated with the dosage effects of X-linked genes, especially in the context of hPGCLCs specification.

Notably, *Sox17* is also transiently expressed in mouse PGCs at E7.5 (Figure 6G).^48^ *Prdm14*, a crucial regulator of mouse PGCs, promotes epigenetic reprogramming and pluripotency.^67^ Overexpression of *Prdm14* alone, but not *Blimp1* or *Tfap2c,* is sufficient to induce the mouse PGC fate *in vitro*.^68^ Sox2 is a prominent target of Prdm14 in the maintenance of pluripotency.^67^ Consequently, Sox17 is diminished as co-existence of Sox2 and Sox17 could be detrimental to cellular viability as demonstrated here. Furthermore, the molecular network involved in PGCs specification has diverged between mice and humans; for instance, PRDM14 targets show a lack of conservation and are vastly different.^69^ In humans, upregulation of SOX17, BLIMP1, and TFAP2C precedes PRDM14 expression.^69^ Thus, abnormally elevated SOX2 expression during hPGCLCs specification could skew the fate towards the ectoderm lineage when PRDM14 is not activated to repress this alternative fate, consistent with our findings.

The molecular mechanisms of cell fate determination, particularly those regulating gene expression, are fundamental to the development and evolution of all organisms. Our findings suggest that the mutual exclusivity of gene expression is essential for cell lineage determination and co-expression of SOX2 and SOX17 is detrimental. SOX2 is an intrinsic determinant for neuroectoderm lineage after gastrulation.^41, 43^ We now found that SOX2 is a downstream effector of X-linked genes in human, which, together with late expression of PRDM14 to repress alternative fate, necessitates SOX17 to specify the PGC fate and eliminate cells with abnormal expression of SOX2 by engaging them in cell death. Intriguingly, mutually exclusive expression of SOX2 and SOX17 are also observed in TGCTs. SOX2 is repressed in seminomas but activated in embryonic carcinomas, a more malignant TGCT enriched with pluripotency and differentiation pathways.^70^ Mutually exclusive gene expression patterns have been widely observed across cancers and are considered as a strategy for tumor initiation and progression.^71^ Overexpression of SOX2 in TCam-2, the first well-characterized seminoma-derived cell line transforms TCam-2 cells to an embryonal carcinoma-like fate.^58^ It would be of great interest to track the transformative process and examine whether SOX2 expression can be titrated to a level insufficient for the embryonal carcinoma transformation.

### Germline entry and mitochondrial dynamics

In mammals, nuclear-encoded mitochondrial genes are underrepresented on the X chromosome. ^72^ This is postulated as an evolutionary adaptation aimed at mitigating potential imbalances in the transmission of genes that are predominantly inherited from one sex, alongside the maternally inherited mitochondria pattern.^72^ Our study unveils an additional nuanced perspective, proposing that X-linked genes may directly or indirectly modulate mitochondrial morphology and function. In this case, the XCI escape gene *USP9X* is implicated in enhancing *SOX2* gene expression and protein stability, thereby regulating mitochondrial morphological dynamics and oxidative phosphorylation function. Germline development intricately intertwines with metabolic shifts and mitochondrial dynamics, characterized by a transition from glycolysis to oxidative phosphorylation highlighting the dynamic energy requisites during this phase.^73^ This metabolic shift entails a reduction in mitochondrial DNA copy number and orchestrated mitochondrial dynamics to facilitate mitochondrial bottleneck selection, more pronounced in females.^74^ Such mitochondrial bottleneck selection is crucial to eliminate mitochondria with deleterious mutations, thereby ensuring mitochondrial integrity in germline transmission.^75^ Our findings revealed that SOX2 upregulation in female and KS results in diminished oxidative phosphorylation and increased mitochondrial clustering, which could subsequently hamper stringent mitochondrial regulation in the germline development. We have demonstrated that SOX2 could potentially inhibit relevant nuclear-encoded mitochondrial genes. Additionally, there is evidence showing that SOX2 could also influence STAT3 translocation into the mitochondria, thereby perturbing mitochondrial DNA-encoded gene expression,^76^ a phenomenon observed in both KS and female lines.

In summary, our study advances our understanding of the detrimental implications of SOX2 during human germ cell development. These findings have established an unprecedented association within X-linked gene dosage effects, germ cell specification, and mitochondrial function. Furthermore, this knowledge holds promise for its application in the *in vitro* culture of hPGCLCs derived from PSCs with an unfavourable XCI states.

## EXPERIMENTAL PROCEDURES

### Cell culture and iMeLCs induction

Human PSCs including iPSCs and ESCs were cultured in NutriStem® hPSC XF Medium (Sartorius, 05-100-1A) after coating with 50 μg/μL Vitronectin (VTN-N) recombinant human protein (Gibco, A14700). PSCs were passed every three days. For iMeLC induction, PSCs were digested by Tryple Select Enzyme (Gibco,12563-029) for 7 minutes in 37 °C incubator after DPBS wash. Cells were centrifuged for 5 min at 300 × g after pipetted into single cells. After cell counting, 60,000 ∼80,000 cells were seeded in each well of fibronectin (Millipore, FC010) coated 12-well plate. IMeLCs were cultured in Glasgow’s MEM medium (Gibco, 11710-035) supplemented with 15% Knockout serum replacement (Gibco, 10828-028), non-essential amino acids solution (Gibco,11140-050), sodium pyruvate (Gibco,11360-070), Penicillin-streptomycin-Glutamine (Gibco, 10378-016), 2-Mercaptoethanol (Gibco, 21985-023), animal-free recombinant Activin A (Pepro tech, AF-120-14E), CHIR99021 (Sigma, SML1046), and Y-27632 (Dihydrochloride) ROCK pathway inhibitor (Stem cell technologies,72307). Culture media were changed every day for PSCs and iMeLCs.

### hPGCLC differentiation

PGCLC differentiation was performed 46∼48 hours after iMeLC induction. IMeLCs were washed with DPBS and digested into single cells by 7 minutes treatment of TrypLE Select Enzyme. After cell counting, 200 μL PGCLC medium containing 5 000 live cells were put into each well of ultra-low attachment 96-well plate. The cells were cultured in 37 °C incubator for 4∼6 days.

The PGCLC medium is 15% Knockout serum replacement (Gibco, 10828-028) Glasgow’s MEM medium (Gibco, 11710-035) with addition of non-essential amino acids solution (Gibco,11140-050), sodium pyruvate (Gibco,11360-070), Penicillin-streptomycin-Glutamine (Gibco, 10378-016), 2-Mercaptoethanol (Gibco, 21985-023), animal-free recombinant Activin A (Pepro tech, AF-120-14E), and cytokine including hLIF (Stem cell, 78055), hEGF (R&D systems, 236-EG-200), hSCF (Peprotech, 300-07), BMP4 (R&D systems, 314-BP-500), Y-27632 (Dihydrochloride) ROCK pathway inhibitor (Stem cell technologies,72307).

### FACS identification of hPGCLC

At 4- and 6- days post differentiation, the efficiency of hPGCLCs were determined by FACS using surface markers EpCAM (Biolegend, 324210) and CD49f (INTEGRINα6, Biolegend, 313624) as previously reported.^24^ Briefly, PGCLC spheres were collected from 96-well plate with truncated pipette. Spheres were dissociated by 8∼10 minutes of TrypLE Select Enzyme treatment after washed by DPBS. After centrifuge, dissociated cells were resuspended with 100 μL FACS buffer containing Alexa Fluor 488 anti-human CD326 (EPCAM) and Brilliant violet 421 anti-human /mouse CD49f (Intergrin- α 6 or IGTA6) on ice for 25∼30 minutes. Centrifuging 5 min at 300 × g after adding 1mL FACS buffer, the cells were resuspended in 400 μL FACS buffer and filtered by 35 μm cell strainer. FACS analyses were performed on BD FACS CANTO II machine.

### Mitochondrial staining

Cells were seeded on coverslip at proper density and stained with MitoTracker™ Red CMXRos (Invitrogen, M7512) or MitoTracker™ Orange CMTMRos (Invitrogen, M7510) for 30 min at 37 °C incubator. Commence with two washes using PBS, followed by fixation in 4% PFA for 10 minutes. After fixation, we did another two PBS washes before incubation in 0.5% Triton X-100 for 10 minutes. Next, we performed two additional washes with PBS and performed DAPI staining for 10 minutes at room temperature. Then we washed three times with PBS. Next, we placed the coverslip on the slide using mounting medium and seal it with nail polish. Finally, we stored the slide at 4 ℃ in a dark environment. The coverslip sample can be used for immunostaining at step just before DAPI staining.

### Immunofluorescence staining

Cells were seeded on coverslip coated with vitronectin or fibronectin. After 2- or 3-days culture, the cells were fixed by 4% PFA for 10 minutes at room temperature after washed with PBS twice. Then, cells were permeabilized with 0.5% Triton X-100 in PBS for 10 min and washed by PBS and incubated with 2% BSA/PBS at room temperature for 1 hour. Next, cells were incubated with primary antibody in 2% BSA/PBS overnight at 4 ℃. Then we washed three times with PBS. The cells were incubated with secondary antibody for 1 h at room temperature and washed three times with PBS., We did DAPI staining for 10 minutes and washed with PBS for three times, 5 minutes each time. Next, we added one drop of mounting medium on slide and put on the coverslip, seal with nail polish and store in dark −20 ℃.

### siRNA treatment

Non-targeting siRNAs control (Scramble controls) and SOX2/USP9X/IGSF1/CHRDL1 siRNA were transfected into cells by using Lipofectamin RNAiMAX Transfection Reagent (Invitrogen, 13778075). Briefly, lipofectamine RNAiMAX and siRNA were diluted by optiMEM medium separately, then add diluted siRNA into diluted Lipofectamin RNAiMAX at 1:1 ratio, gently mix, incubator for 5 minutes at room temperature, then add siRNA-lipid complex into cells. Change it to a fresh medium after 4∼6 hours.

### Primer-seq

Primer-seq library preparation was performed as previously described^77^ and following protocol https://www.protocols.io/view/prime-seq-81wgb1pw3vpk/v2/materials. Briefly, 40 ng RNA was reverse transcribed into cDNA and barcoded with oligo dT (E3V7NEXT) by PCR thermocycler, then cleaned with 22% PEG beads and eluted in 17 μL H_2_O. After removing ssDNA and primers by Exonuclease I, the cleaned cDNA was pre-amplified by KAPA HIFI 2 × enzyme. The beads were purified with 22 PEG beads and eluted with 10 μL H_2_O. The amplified dsDNA was fragmented, end repaired and dA-tailing by Ultra II FS Enzyme. Then ligation was performed with primer-seq adapter. Cleaned with SRPI select beads and eluted with 11 μL 0.1×TE, the resultant cDNA was ligated with Nextera i7 and Truseq i5 index primer. Finally, the cDNA was amplified with Q5 Master Mix (M0544L) and sequenced pair-end 150 base pairs on Novaseq 6000 platform after SRPI clean.

### Seahorse

Agilent seahorse XF96 cell culture microplate (Agilent technologies,101085-004) was first coated with 30 μL poly-D-lysine (P6407, 50 μg/mL) for 1 hour at room temperature, washed with 200 μL H_2_O and air dried. Then the microplate was coated with 50 μL vitronectin (50 μg/μL) for 1 hour at room temperature. 10^4^ PSCs in 50 μL NutriStem medium plus ROCKi were seeded into each well after coating. We left the well of each corner empty on the plate. At 24 hours after seeding, cells were sequentially treated with Oligomycin, FCCP, Antimycin/rotenone and XF media. Seahorse was run on Agilent seahorse XF96 machine.

### Real-time oxygen consumption measurement

PSCs, iMeLCs and hPGCLC were cultured in corresponding 96-well plate independently, with 5000 live cells seeded in each well. The O_2_ consumption was monitored every minute by a RESIPHER device (Lucid Scientific) with probes on the 96-well plate lid. For PSCs and iMeLCs, culture medium was changed every 24 hours. Cell numbers were counted manually by cytometer using parallel 96-well group cultured cells when changing medium. Each group has 4 well of 96-well replicates. Finally, the O_2_ consumption is normalized by cell number at around 24 or 48 hours after seeding.

### Prime-seq alignment and processing

For the processing of Prime-seq data, quality control checks were conducted using FastQC (v0.11.8) to assess the raw sequence data. Subsequent to quality inspection, poly(A) tails were excised using Cutadapt (v1.14).^78^ The prepped sequences underwent further processing with the zUMIs^79^ pipeline (v2.9.4), which included filtering, alignment, and quantification steps, applying a Phred quality score cutoff of 20 for 2 barcode bases and 3 UMI bases. Mapping of the curated sequences was performed against the human reference genome (GRCh38) with Gencode annotations (v35) via the STAR aligner (v2.7.3a).^80^ For the identification of differentially expressed genes, EdgeR was employed, setting a threshold for significance at an FDR of less than 0.05 and a fold change greater than 1.5. Gene set enrichment analysis was performed with Genekitr.^81^

### Single-cell alignment and processing

For the processing of Smart-seq3 data, sequencing reads were aligned to the human genome (hg38) and quantified using the zUMIs pipeline,^79^ following the default settings tailored for Smart-seq3, as previously documented.^82^ For subsequent analyses, Seurat V4^83^ was the primary tool unless specified otherwise.

The initial step involved quality control measures to exclude cells with a low number of genes and counts, as determined by Seurat’s violin plots. Additionally, cells exhibiting more than 10% mitochondrial content were removed. Following this, the expression matrix underwent Log normalization and scaling using Seurat’s default parameters. Dimensional reduction was carried out using PCA, focusing on 3000 variable genes. The selection of principal components (PCs) and harmony dimensions for cell clustering and UMAP (Uniform Manifold Approximation and Projection) visualization was guided by an Elbow plot, which aids in identifying a significant inflection point in the data. In silico bulk RNA-seq data were generated by aggregating the single-cell expression profiles with Seurat V4. Slingshot^84^ was employed for pseudotime trajectory analysis. Unlike other analyses, the data for Slingshot^84^ were not scaled by variance as genes contribute differently in trajectory analysis. Differentially expressed genes between groups or clusters were identified using MAST^85^ with a criterion of log2(fold change) greater than 1 and an adjusted p-value below 0.05. Temporal gene expressions along pseudotime were modeled using TradeSeq.^86^ Quadratic programming for cell fate identification was performed using R package quadprog, following the same procedure as previously reported.^44^ The hES, ectoderm and mesoendoderm gene expression reference were obtained from previous study.^45^

### Expression ratios

For the calculation of autosomal expression ratios, we used protocol as previously reported.^87^ First, we focused on genes exhibiting expression levels greater than 1 TPM. These ratios were determined relative to the median expression of autosomes, specifically excluding genes known to escape X-Chromosome Inactivation (XCI). To address the variance in gene numbers across different chromosomes, a bootstrapping approach was employed. In this method, random sets of autosomal genes, matching the size of the chromosome under investigation, were selected to serve as a comparative background. This process was replicated 1,000 times (n = 1,000) to ensure statistical robustness and accuracy in the ratio calculations.

### ChIP-seq data analysis

Previously published ChIP-Seq peak bed and bigwig data were used. Heatmap of the peak intensities were visualized using the bigwig data with Deeptools2.^88^ Motif enrichment was analyzed with findmotifs.pl from Homer.^47^

### Data availability

The human sequencing data will be uploaded to human genome archive with controlled access. This paper also included public data. PSC SOX2 ChIP-Seq is from GSE81899; hPGCLC SOX17 ChIP-seq is from GSE159654.

## AUTHOR CONTRIBUTIONS

Conceptualization, W.H., Q.L., J.Z. and Q.D.; methodology, W.H., Q.L., J.Z. and Q.D.; investigation, W.H., Q.L., J.Z. and Q.D with assistance from A.Z., M.W., L.F., A.Z., A.R., E.L, J.S., J.C.; formal analysis: W.H., Q.L., J.Z. and Q.D..; writing, W.H., Q.L., J.Z. and Q.D..; review and editing, all authors.

## ACKNOWLEDGMENTS

We thank Hannah Schorle for technical assistance for Seahorse analysis. Q.D. is Wallenberg Academy Fellow in Medicine and is also supported by the Swedish Research Council (no. 2018-02557 and 2020-00253), Åke Wibergs Stiftelse, Birgitta and Carl-Axel Rydbeck’s Research grant for Pediatric Research (no. 2022-00327), and faculty funding at Karolinska Institutet. Q.L. is supported by Chinese Scholarship Council.

