## Supplemental Figure S1-S6 for "Divergent roles of SOX2 in human and mouse germ cell specification related to X-linked gene dosage effects"

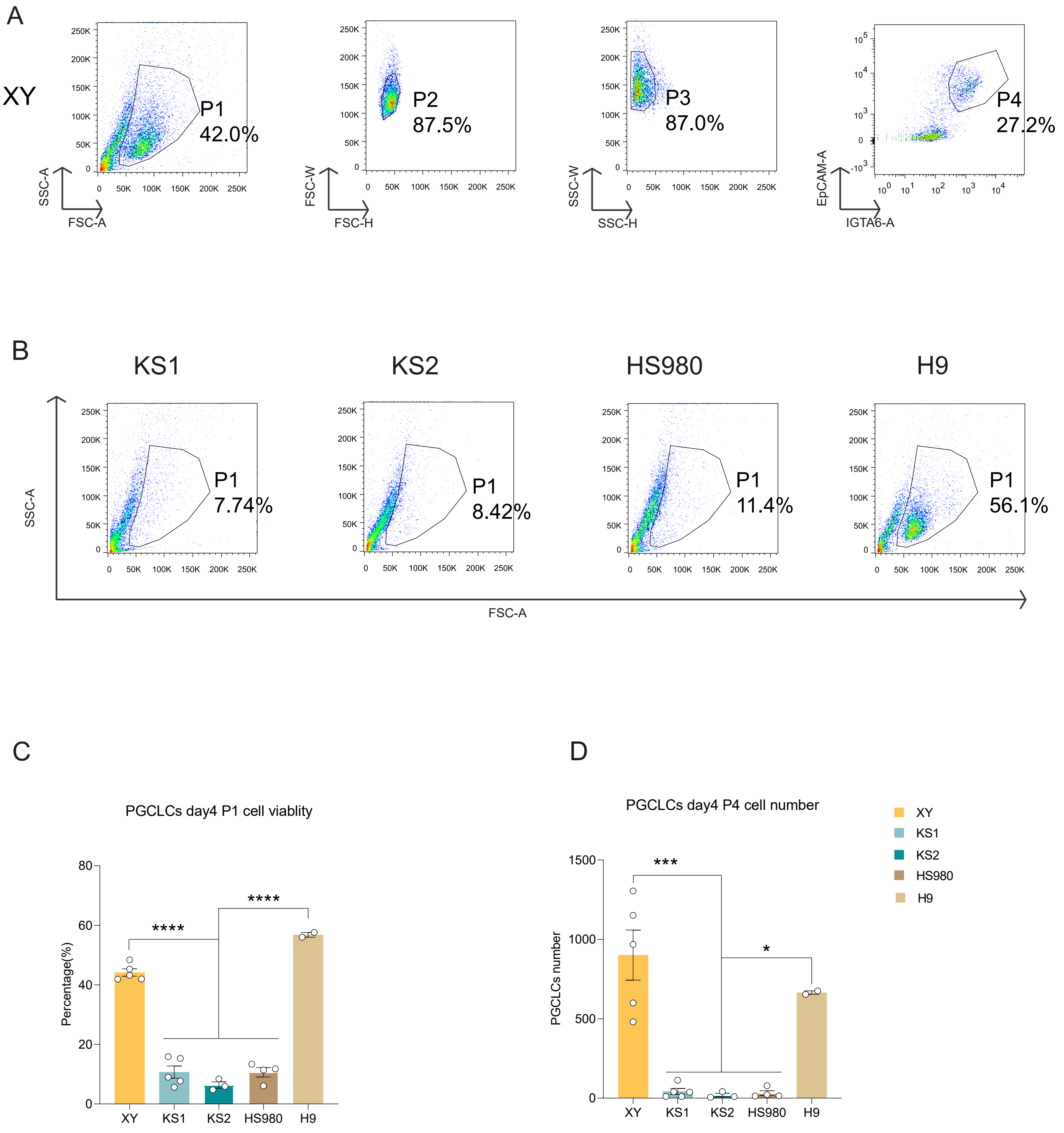

A
XY
P1
42.0%
P2
87.5%
P3
87.0%
P4
27.2%
SSC-A
FSC-A
FSC-W
FSC-H
SSC-W
SSC-H
EpCAM-A
IGTA6-A
B
KS1
KS2
HS980
H9
P1
7.74%
P1
8.42%
P1
11.4%
P1
56.1%
SSC-A
FSC-A
C
PGCLCs day4 P1 cell viablity
Percentage(%)
****
****
XY
KS1
KS2
HS980
H9
D
PGCLCs day4 P4 cell number
PGCLCs number
***
*
XY
KS1
KS2
HS980
H9

**SUPPLEMENTAL FIGURE LEGENDS**

**Figure S1. X-linked gene dosage underlies the differentiation efficiency of hPGCLCs, Related to Figure 1.**

(A) FACS analysis of XY cells from P1 to P4. P1 represents living cell population, P2 and P3 is to gate single cell population for analysis, in the end, P4 represents the PGCLCs identified by EpCAM and IGTA6.

(B) FACS analysis of P1 cells in KS1, KS2, HS980 and H9.

(C) Statistical analysis of P1 viable cells in PGCLCs day 4 of five cell lines.

(D) Statistical analysis of P4 cell number in PGCLCs day 4 of five cell lines.

Data in (C), (D) are shown as mean ± SEM. ****$p < 0.0001$ by one-way ANOVA comparison.

Figure S2

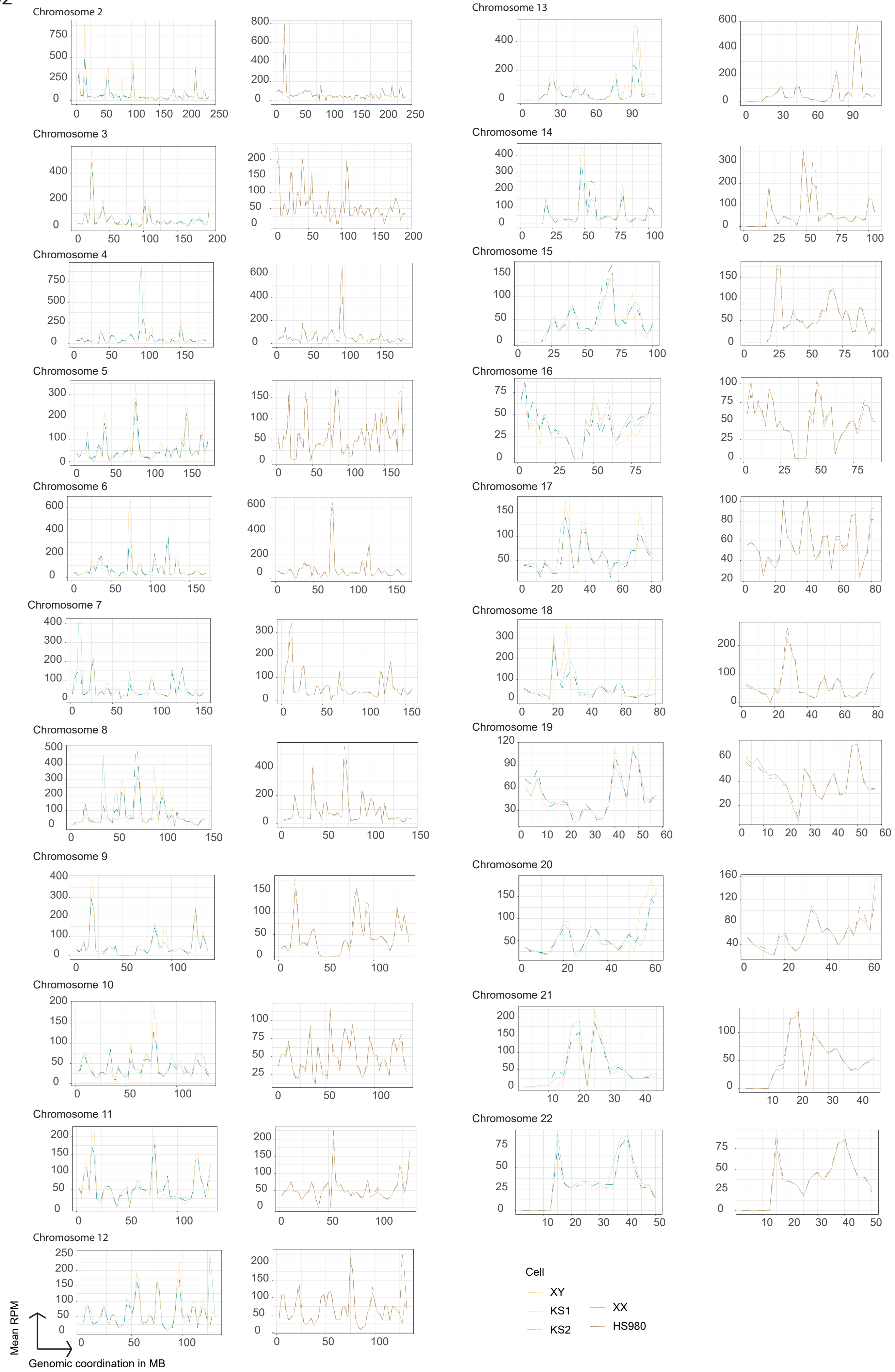

**Figure S2. Autosomal gene expressions on genomic coordination, Related to Figure 1.**

Figure S3

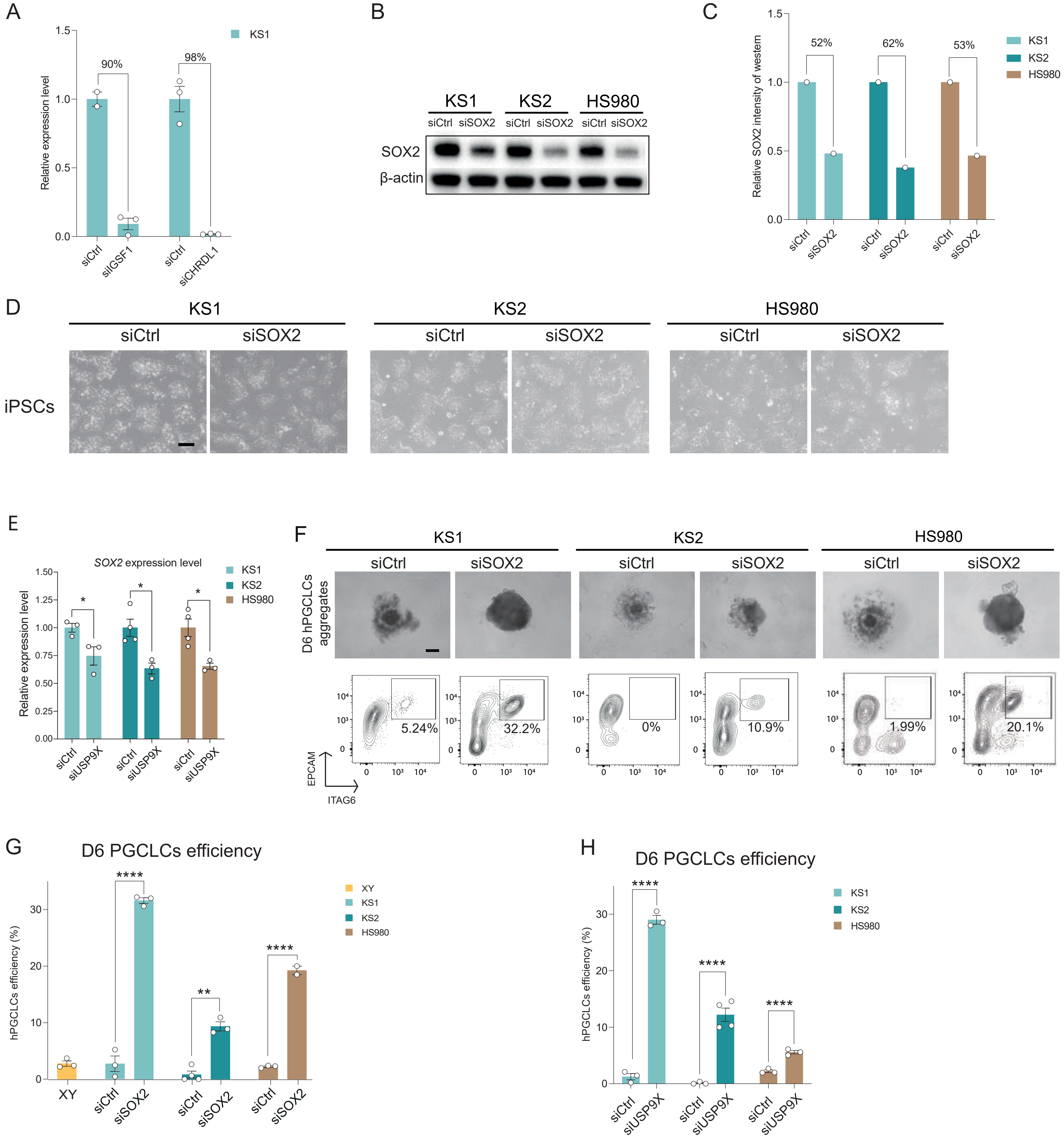
A
KS1
Relative expression level
90%
98%
siCtrl
siIGSF1
siCtrl
siCHRDL1
B
KS1
KS2
HS980
siCtrl
siSOX2
SOX2
β-actin
C
Relative SOX2 intensity of western
52%
62%
53%
KS1
KS2
HS980
D
iPSCs
E
SOX2 expression level
Relative expression level
siUSP9X
F
D6 hPGCLCs aggregates
5.24%
32.2%
0%
10.9%
1.99%
20.1%
EPCAM
ITAG6
G
D6 PGCLCs efficiency
hPGCLCs efficiency (%)
XY
H

**Figure S3. Knockdown efficiencies, Related to Figure 3.**
(A) Knockdown efficiency of *IGSF1* and *CHRDL1* in KS1 iPSCs.
(B) Western blotting showing the decrease of SOX2 protein levels in KS1, KS2, and HS980 after SOX2 KD.
(C) Quantitative analysis of western blotting in B.
(D) Representative image of the PSC morphology of KS1, KS2, and HS980 after 72 hours of SOX2 knockdown. Scale bar is 200 μm.
(E) SOX2 expression level after *USP9X* KD by real-time qPCR.
(F) D6 hPGCLC spheroid and FACS identification of hPGCLC after SOX2 KD in KS1, KS2 and HS980, respectively. Scale bar, 200 μm.
(G) Statistics of D6 hPGCLC efficiencies after SOX2 KD in KS1, KS2 and HS980 from three independent experiments.
**(**H**)** Statistics of D6 hPGCLC efficiencies after USP9X KD in KS1, KS2 and HS980 from three independent experiments.
Data in (A), (E), (G), (H) are shown as mean ± SEM. n.s., non-significant; **p* < 0.05; ***p* < 0.01; *****p* < 0.0001 by t test.

Figure S4

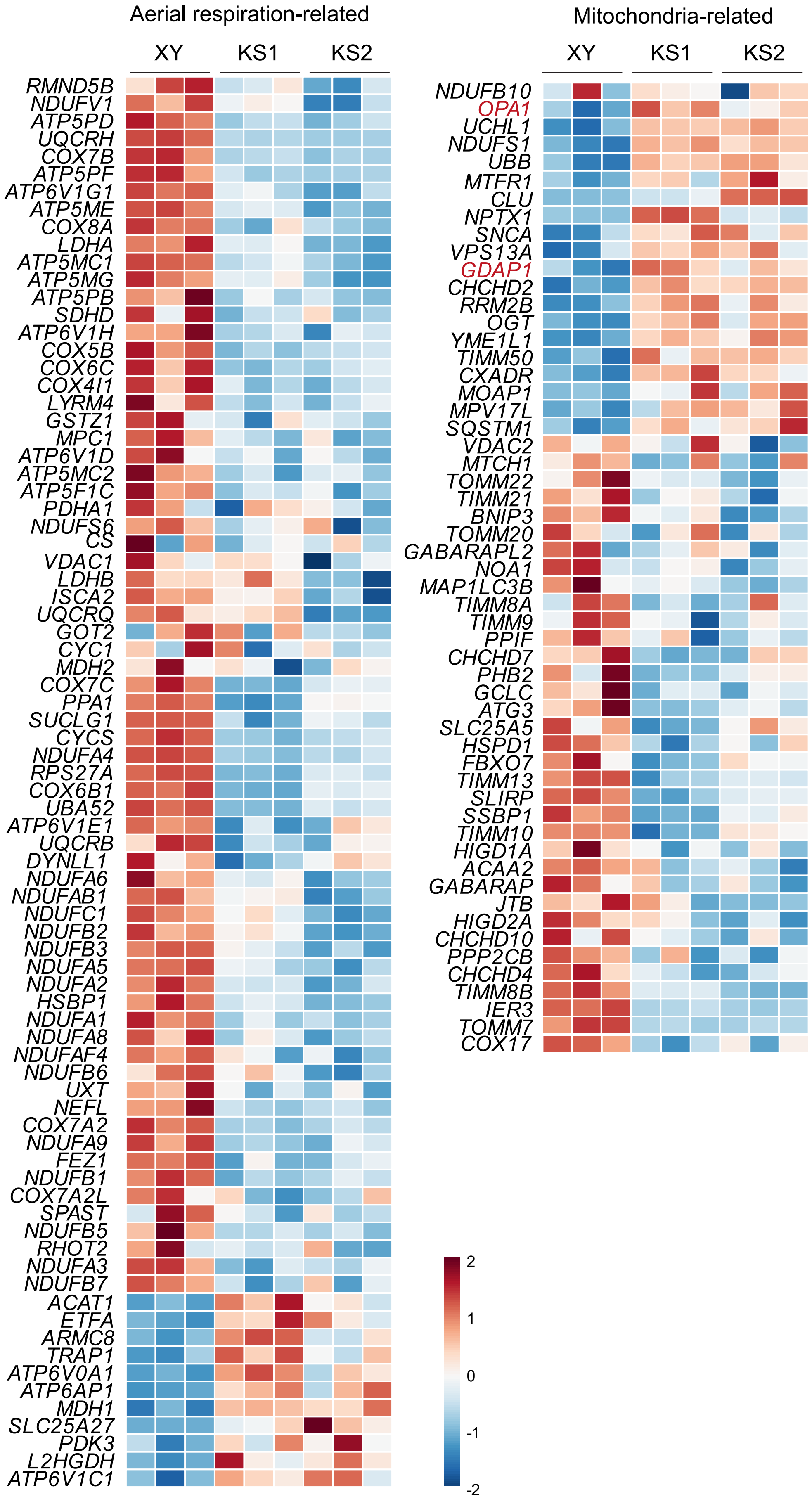

**Figure S4. Heatmap showing the KS common DEGs related with mitochondria and aerobic respiration pathways, Related to Figure 4.**

Figure S5

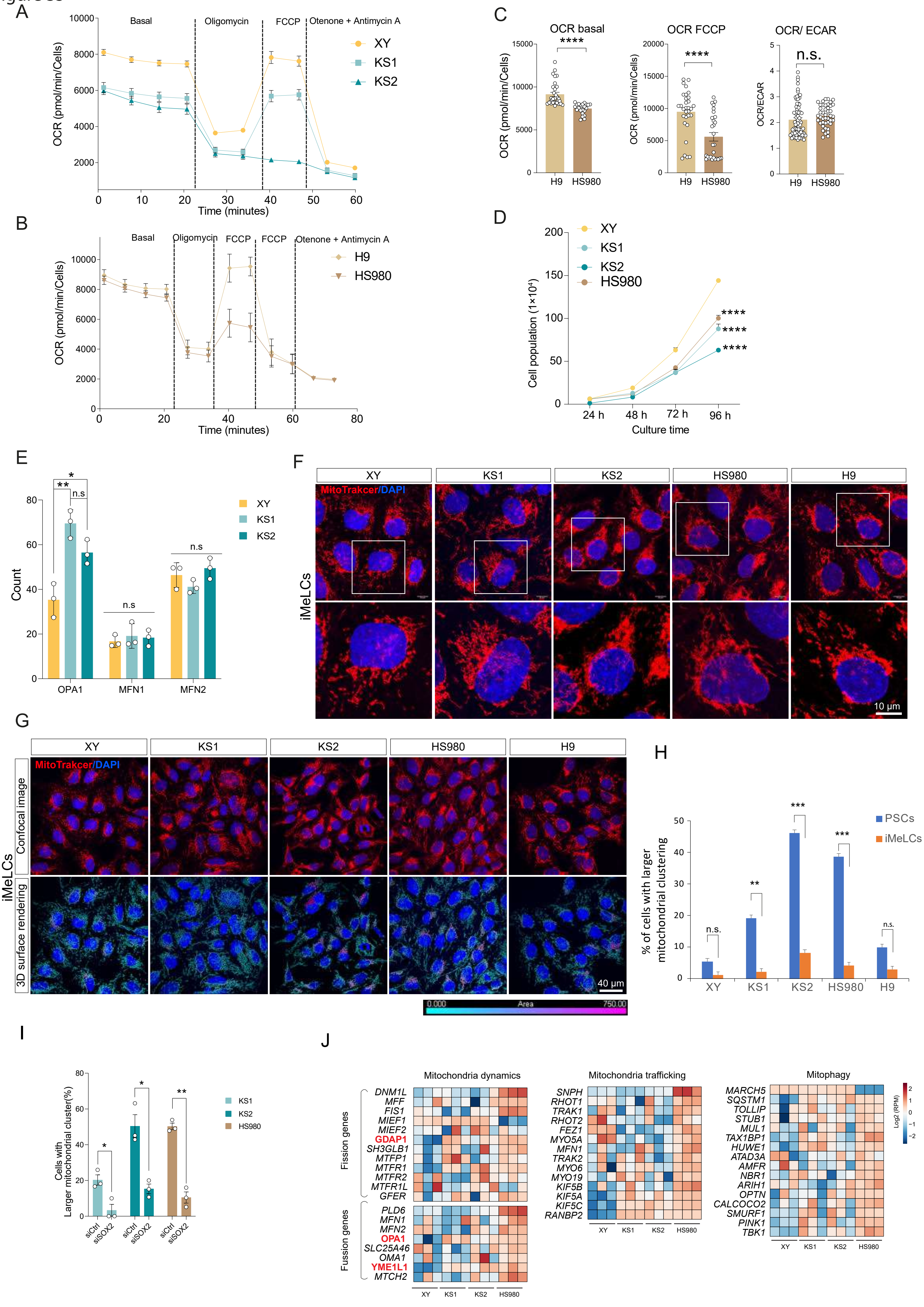

A
Basal
Oligomycin
FCCP
Otenone + Antimycin A
XY
KS1
KS2
OCR (pmol/min/Cells)
Time (minutes)
B
Basal
Oligomycin
FCCP
FCCP
Otenone + Antimycin A
H9
HS980
OCR (pmol/min/Cells)
Time (minutes)
C
OCR basal
OCR FCCP
OCR/ ECAR
****
****
n.s.
OCR (pmol/min/Cells)
OCR/ECAR
H9
HS980
D
XY
KS1
KS2
HS980
Cell population (1×10⁴)
24 h
48 h
72 h
96 h
Culture time
E
Count
OPA1
MFN1
MFN2
n.s
XY
KS1
KS2
F
XY
KS1
KS2
HS980
H9
MitoTrakcer/DAPI
iMeLCs
10 μm
G
XY
KS1
KS2
HS980
H9
MitoTrakcer/DAPI
Confocal image
3D surface rendering
iMeLCs
40 μm
0.000
Area
750.00
H
PSCs
iMeLCs
% of cells with larger mitochondrial clustering
n.s.
**
***
***
n.s.
XY
KS1
KS2
HS980
H9
I
Cells with Larger mitochondrial cluster(%)
KS1
KS2
HS980
siCtrl
siSOX2
J
Mitochondria dynamics
Fission genes
DNM1L
MFF
FIS1
MIEF1
MIEF2
GDAP1
SH3GLB1
MTFP1
MTFR1
MTFR2
MTFR1L
GFER
Fusion genes
PLD6
MFN1
MFN2
OPA1
SLC25A46
OMA1
YME1L1
MTCH2
Mitochondria trafficking
SNPH
RHOT1
TRAK1
RHOT2
FEZ1
MYO5A
MFN1
TRAK2
MYO6
MYO19
KIF5B
KIF5A
KIF5C
RANBP2
Mitophagy
MARCH5
SQSTM1
TOLLIP
STUB1
MUL1
TAX1BP1
HUWE1
ATAD3A
AMFR
NBR1
ARIH1
OPTN
CALCOCO2
SMURF1
PINK1
TBK1
XY
KS1
KS2
HS980
Log2 (RPM)

**Figure S5. SOX2 reduces cellular oxidative phosphorylation and influences mitochondria morphology, Related to Figure 4.**
(A) Seahorse data of the XY, KS1, and KS2 PSCs.
(B) Seahorse data of the HS980, and H9 PSCs.
(C) Statistical analysis OCR of basal and FCCP in B.
(D) Growth curve of the PSCs of XY, KS1, KS2, and HS980.
(E) OPA1, MFN1, MFN2 expression level analysis of XY, KS1, KS2 iPSCs from bulk RNA sequenced data.
(F) Mitochondrial morphology of the iMeLCs of XY, KS1, KS2 and HS980 iMeLCs from confocal microscope.
(G) 3D mitochondrial morphology and large cluster in the iMeLCs of XY, KS1, KS2 and HS980 from Imaris software analysis.
(H) Percentage of cells with mitochondria clusters in the iMeLCs XY, KS1, KS2 and HS980. Cluster is designated as surface area> 500 $\mu m^2$.
(I) Percentage of cells with mitochondria clusters in the iPSCs XY, KS1, and KS2 after SOX2 KD. Cluster is designated as surface area> 500 $\mu m^2$.
(J) Heatmaps showing the expressions of selected mitochondria related genes.
Data in (C), (E), (H), (I) are shown as mean ± SEM. n.s., non-significant; $*p < 0.05$; $**p < 0.01$; $****p < 0.0001$ by t test.
Data in (D) are shown as mean ± SEM. $****p < 0.0001$ by two-way ANOVA comparison.

Figure S6

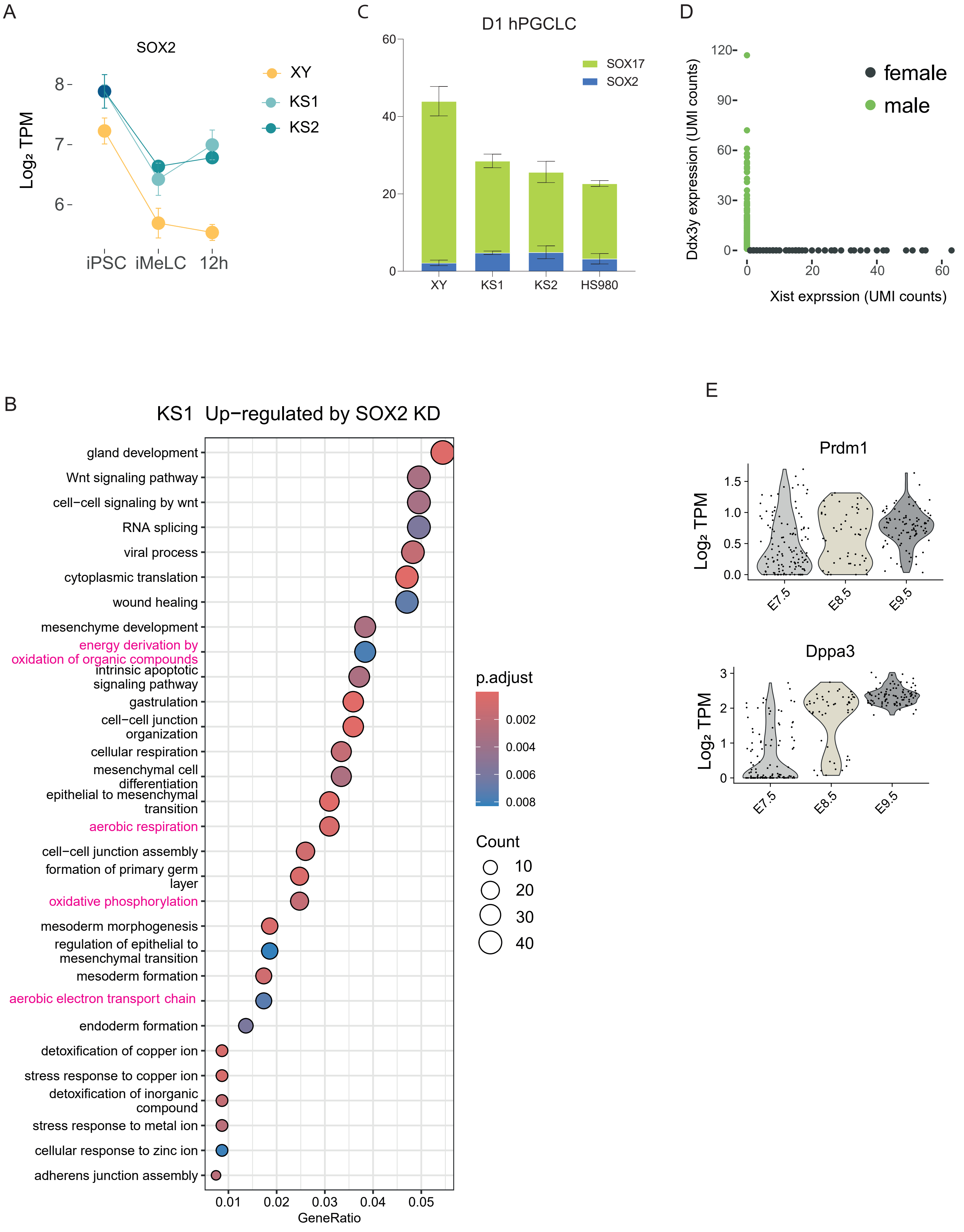
A
SOX2
Log₂ TPM
8
7
6
iPSC
iMeLC
12h
XY
KS1
KS2
C
D1 hPGCLC
60
40
20
0
SOX17
SOX2
XY
KS1
KS2
HS980
D
Ddx3y expression (UMI counts)
120
90
60
30
0
0
20
40
60
Xist exprssion (UMI counts)
female
male
B
KS1 Up−regulated by SOX2 KD
gland development
Wnt signaling pathway
cell−cell signaling by wnt
RNA splicing
viral process
cytoplasmic translation
wound healing
mesenchyme development
energy derivation by oxidation of organic compounds
intrinsic apoptotic signaling pathway
gastrulation
cell−cell junction organization
cellular respiration
mesenchymal cell differentiation
epithelial to mesenchymal transition
aerobic respiration
cell−cell junction assembly
formation of primary germ layer
oxidative phosphorylation
mesoderm morphogenesis
regulation of epithelial to mesenchymal transition
mesoderm formation
aerobic electron transport chain
endoderm formation
detoxification of copper ion
stress response to copper ion
detoxification of inorganic compound
stress response to metal ion
cellular response to zinc ion
adherens junction assembly
0.01
0.02
0.03
0.04
0.05
GeneRatio
p.adjust
0.002
0.004
0.006
0.008
Count
10
20
30
40
E
Prdm1
Log₂ TPM
1.5
1.0
0.5
0.0
E7.5
E8.5
E9.5
Dppa3
Log₂ TPM
3
2
1
0
E7.5
E8.5
E9.5

**Figure S6. mPGC sex related and PGC marker gene expressions, Related to Figure 5 and 6.**

(A) SOX2 gene expression changes in the iMeLC and 12h post BMP induction stages in KS1, KS2 and XY.

(B) Up regulated pathways after SOX2 KD in the 12h stage of KS1.

(C) Statistics of SOX2- and SOX17-expressing cell in D4 hPGCLC spheroids of XY, KS1, KS2, and HS980.

(D) Exclusive gene expressions of *Xist* and *Ddx3y* showing female and male PGCs.

(E) Expression of *Prdm1* and *Dppa3* in mPGCs.

**SUPPLEMENTAL INFORMATION**

KEY RESOURCES TABLE

| REAGENT or RESOURCE | SOURCE | IDENTIFIER |
|---|---|---|
| Antibodies | | |
| Human/Mouse/Rat SOX2 Antibody | R&D systems | MAB2018 |
| Human NAONG antibody | Miltenyi Biotec | 130-104-732 |
| Alexa Fluor® 488 anti-human CD326 (EpCAM) antibody | Biolegend | 324210 |
| Brilliant Violet 421™ anti-human/mouse CD49f (Integrin-a6 or ITGA6) Antibody | Biolegend, | 313624 |
| Chemicals, peptides, and recombinant proteins | | |
| TRI Reagent | Sigma | T9424 |
| Vitronectin (VTN-N) Recombinant Human Protein, Truncated | Fisher Scientific | A14700 |
| Human plasma fibronectin purified protein | Millipore Sigma | FC010 |
| Recombinant Human BMP-4 Protein | R&D Systems | 314-BP |
| Recombinant Human EGF Protein | R&D Systems | 236-EG |
| Recombinant Human SCF | PeproTech | 300-07 |
| Human Recombinant LIF | STEMCELL Technologies | 78055 |
| Y-27632 (Dihydrochloride) | STEMCELL Technologies | 72307 |
| Animal-Free Recombinant Human/Murine/Rat Activin A | PeproTech | AF-120-14E |
| Knockout™ Serum Replacement | Gibco | 10828-028 |
| NutriStem® hPSC XF Medium | Sartorius | 05-100-1A |
| RPMI 1640 Medium | Sigma-Aldrich | R8758 |
| Glasgow's MEM (GMEM) | Thermo Fisher Scientific | 11710035 |
| 2-Mercaptoethanol | Thermo Scientific | 21985023 |
| Sodium pyruvate | Thermo Fisher Scientific | 11360–070 |
| TrypLE™ Select Enzyme (1X), no phenol red | Thermo Scientific | 12563029 |
| MEM α, nucleosides, GlutaMAX™ Supplemen | Thermo Scientific | 32571036 |
| DPBS, no calcium, no magnesium | Thermo Fisher Scientific | 14190094 |
| Poly-D-lysine hydrobromide | Sigma | P6407 |
| Doxycycline hyclate | Merck | 33429 |
| Critical commercial assays | | |

| | | |
|---|---|---|
| Lipofectamine RNAiMAX Transfection Reagent | Fisher Scientific | 13-778-075 |
| Applied Biosystems Power UP SYBR Green PCR Master Mix | Fisher Scientific | 15390929 |
| 5X All-In-One RT MasterMix | ABM | G490 |
| QIAGEN Plasmid Plus Midi Kit | Qiagen | 12943 |
| Experimental models: Cell lines | | |
| HS980 | Rodin et al., (2014) | N/A |
| XY | Panula, S., et al. (2019) | N/A |
| KS1 | Panula, S., et al. (2019) | N/A |
| KS2 | Panula, S., et al. (2019) | N/A |
| Oligonucleotides | | |
| Control siRNA | | |
| ON-TARGETplus Human SOX2 (6657) siRNA - SMARTpool | Horizon Discovery | L-011778-00-0005 |
| USP9X siRNA | ThermoFisher Scientific | 105099 |
| CHRDL1 siRNA | ThermoFisher Scientific | S534487/s40796 |
| IGSF1 siRNA | ThermoFisher Scientific | s535148/s230594 |
| Recombinant DNA | | |
| Software and algorithms | | |
